# Differential global distribution of marine picocyanobacteria gene clusters reveals distinct niche-related adaptive strategies

**DOI:** 10.1101/2022.08.17.504260

**Authors:** Hugo Doré, Ulysse Guyet, Jade Leconte, Gregory K. Farrant, Benjamin Alric, Morgane Ratin, Martin Ostrowski, Mathilde Ferrieux, Loraine Brillet-Guéguen, Mark Hoebeke, Jukka Siltanen, Gildas Le Corguillé, Erwan Corre, Patrick Wincker, David J. Scanlan, Damien Eveillard, Frédéric Partensky, Laurence Garczarek

## Abstract

The ever-increasing number of available microbial genomes and metagenomes provide new opportunities to investigate the links between niche partitioning and genome evolution in the ocean, notably for the abundant and ubiquitous marine picocyanobacteria *Prochlorococcus* and *Synechococcus*. Here, by combining metagenome analyses of the *Tara* Oceans dataset with comparative genomics, including phyletic patterns and genomic context of individual genes from 256 reference genomes, we first showed that picocyanobacterial communities thriving in different niches possess distinct gene repertoires. We then managed to identify clusters of adjacent genes that display specific distribution patterns in the field (CAGs) and are thus potentially involved in the adaptation to particular environmental niches. Several CAGs are likely involved in the uptake or incorporation of complex organic forms of nutrients, such as guanidine, cyanate, cyanide, pyrimidine or phosphonates, which might be either directly used by cells, for e.g. the biosynthesis of proteins or DNA, or degraded into inorganic nitrogen and/or phosphorus forms. We also highlight the frequent presence of CAGs involved in polysaccharide capsule biosynthesis in *Synechococcus* populations thriving in both nitrogen- and phosphorus-depleted areas, which are absent in low-iron regions, suggesting that the complexes they encode may be too energy-consuming for picocyanobacteria thriving in these areas. In contrast, *Prochlorococcus* populations thriving in iron-depleted areas specifically possess an alternative respiratory terminal oxidase, potentially involved in the reduction of Fe(III) into Fe(II). Together, this study provides insights into how these key members of the phytoplankton community might behave in response to ongoing global change.

**Significance Statement:** Picocyanobacteria face various environmental conditions in the ocean and numerous studies have shown that genetically distinct ecotypes colonize different niches. Yet the functional basis of their adaptation remains poorly known, essentially due to the large number of genes of yet unknown function, many of which have little or no beneficial effect on fitness. Here, by combining comparative genomics and metagenomics approaches, we have identified not only single genes but also entire gene clusters, potentially involved in niche adaptation. Although being sometimes present in only one or a few sequenced strains, they occur in a large part of the population in specific ecological niches and thus constitute precious targets for elucidating the biochemical function of yet unknown niche-related genes.

## Introduction

During the last two decades the sequencing of a large number of microbial genomes (more than 425,000 were available in Genbank in July 2022) has allowed tremendous advances in the elineation of core, accessory and unique gene repertoires within closely related organisms by building clusters of likely orthologous genes (CLOGs) based on sequence homology (1–4).Although this approach was tentatively used to identify the genetic basis of niche adaptation, relatively few genes were identified as being specific to particular ecotypes and thus potentially involved in niche adaptation (5–9). Various reasons may underpin this difficulty to identify niche-specific genes by a mere comparative genomics approach. These include the still fairly low number of genomes available given extensive known microbial genomic diversity (10), a lack of ecological representation due to cultivation biases, a limited knowledge of physiological traits of equenced strains and/or the imprecise delineation of ecotypes and of the limits of their realized environmental niches *sensu* (11), especially for lineages present in low abundance in the field (12–14).

An alternative to comparative genomics to better decipher the link between niche partitioning and genome evolution consists of using the rapidly growing number of metagenomes. Besides triggering the generation of numerous metagenome-assembled genomes (MAGs), allowing to fill the gap for yet uncultured microbial taxa and/or ecotypes (15, 16), metagenome recruitment analyses using reference genomes have also allowed scientists to identify spatial or temporal niche-specific genes (17–19). In this context, due to their abundance and ubiquity in the field and the numerous available genomes, single amplified genomes (SAGs) and MAGs, marine picocyanobacteria constitute highly pertinent models for these metagenomic recruitment approaches. The *Prochlorococcus* and *Synechococcus* genera are indeed the two most abundant members of the phytoplankton community, *Prochlorococcus* being restricted to the 40°S-50°N latitudinal band, while *Synechococcus* distribution extends from the equator to subpolar waters (20, 21). Furthermore, physiological and environmental studies have allowed scientists to decipher their genetic diversity and their main physiological traits as well as toelineate ecotypes or Ecologically Significant Taxonomic Units (ESTUs), i.e., genetic groups within clades occupying a given ecological niche, notably using *Tara* oceans metagenomic data at the global scale (22). While three major ESTU assemblages were identified for *Prochlorococcus* in surface waters, whose distribution was found to be mainly driven by temperature and iron (Fe) availability, eight distinct assemblages were identified for *Synechococcus* depending on three main environmental parameters (temperature, Fe and phosphate availability). Nevertheless, few studies have so far integrated our wide knowledge of ecotype distributions and the genetic and functional diversity of these organisms to identify niche- and/or ecotype-specific genes based on their relative abundance in the field (12, 23–26). Furthermore, most of these previous studies have focused on the abundance of individual genes, or more rarely, on a few genomic regions with known functions, e.g. involved in nitrogen or phosphorus uptake and assimilation (27, 28).

Here, by using a network approach to integrate metagenome analyses of the *Tara* Oceans dataset and synteny of individual accessory genes in 256 reference genomes, MAGs and SAGs, we managed to identify clusters of adjacent genes that display specific distribution patterns along the *Tara* Oceans transect. This led us to the unveil niche- and/or ecotype-specific genomic regions, including several previously unreported and sometimes only present in a few or even single genomes, potentially involved in the adaptation to the main ecological niches occurring in the marine environment (N, P and/or Fe-limited as well as cold vs. warm areas). Delineation of these gene clusters also led us to predict the putative functions of previously uncharacterized genes in these genomic regions based on genes functionally annotated in the same cluster. Altogether, this study provides unique insights into the functional basis of microbial niche partitioning and the molecular bases of fitness in key members of the phytoplankton community.

## Results and Discussion

### Different picocyanobacterial communities exhibit distinct gene repertoires

To analyze the distribution of *Prochlorococcus* and *Synechococcus* reads along the *Tara* Oceans transect, metagenomic reads corresponding to the bacterial size fraction were recruited against 256 picocyanobacterial reference genomes, including 178 whole genome sequences (WGS), and a selection of 48 SAGs and 30 MAGs, primarily representative of still uncultured lineages (e.g. *Prochlorococcus* HLIII-IV, *Synechococcus* EnvA or EnvB). This yielded a total of 1.07 billion recruited reads, of which 87.7% mapped onto *Prochlorococcus* genomes and 12.3% onto *Synechococcus* ones, which were then functionally assigned by mapping them on the manually curated Cyanorak v2.1 CLOG database (29). In order to identify picocyanobacterial genes potentially involved in niche adaptation, we analyzed the distribution across the oceans of flexible genes (i.e., non-core genes in Cyanorak *Prochlorococcus* and *Synechococcus* reference genomes). *Tara* Oceans stations were first clustered according to the relative abundance of flexible genes. This clustering resulted in three well-defined clusters for *Prochlorococcus* (left tree in Fig. 1A), which matched quite well those obtained when stations were clustered according to the relative abundance of *Prochlorococcus* ESTUs, as assessed using the high-resolution marker gene *petB*, encoding the cytochrome *b*_6_ (right tree in Fig. 1A; see also (22)). Only a few discrepancies can be observed between the two trees, including stations *TARA*-070 that displayed one of the most disparate ESTU compositions and *TARA*-094, dominated by the rare HLID ESTU (Fig. 1A). For *Synechococcus*, there was also a good consistency between dendrograms obtained from flexible gene abundance and relative abundance of ESTUs (Fig. 1B). Of the eight assemblages of stations discriminated based on the relative abundance of ESTUs (Fig. 1B), most were retrieved in the clustering based on flexible gene abundance, except for a few intra-assemblage switches between stations, notably those dominated by ESTU IIA (Fig. 1B). Despite these few variations between *Synechococcus* trees, four major clusters can be clearly delineated in both trees, corresponding to four broadly defined ecological niches, namely i) cold, nutrient-rich, pelagic or coastal environments (blue and light red in Fig. 1B), ii) Fe-limited environments (purple and grey), iii) temperate, P-depleted, Fe-replete areas (yellow) and iv) warm, N-depleted, Fe-replete regions (dark red). This correspondence between taxonomic and functional information was also confirmed by the high congruence between distance matrices based on ESTU relative abundance and on CLOG relative abundance (p-value < 10^−4^, mantel test r=0.84 and r=0.75 for *Synechococcus* and *Prochlorococcus*, respectively; dataset 1-4). Altogether, this indicates that distinct picocyanobacterial communities, as assessed based on a single taxonomic marker, also display different gene repertoires. As previously suggested for *Prochlorococcus* (30), this strong correlation between taxonomy and gene content strengthens the idea that, in both genera, the evolution of the accessory genome mainly occurs by vertical transmission, with a relatively low extent of lateral gene transfer.

**Figure 1.**
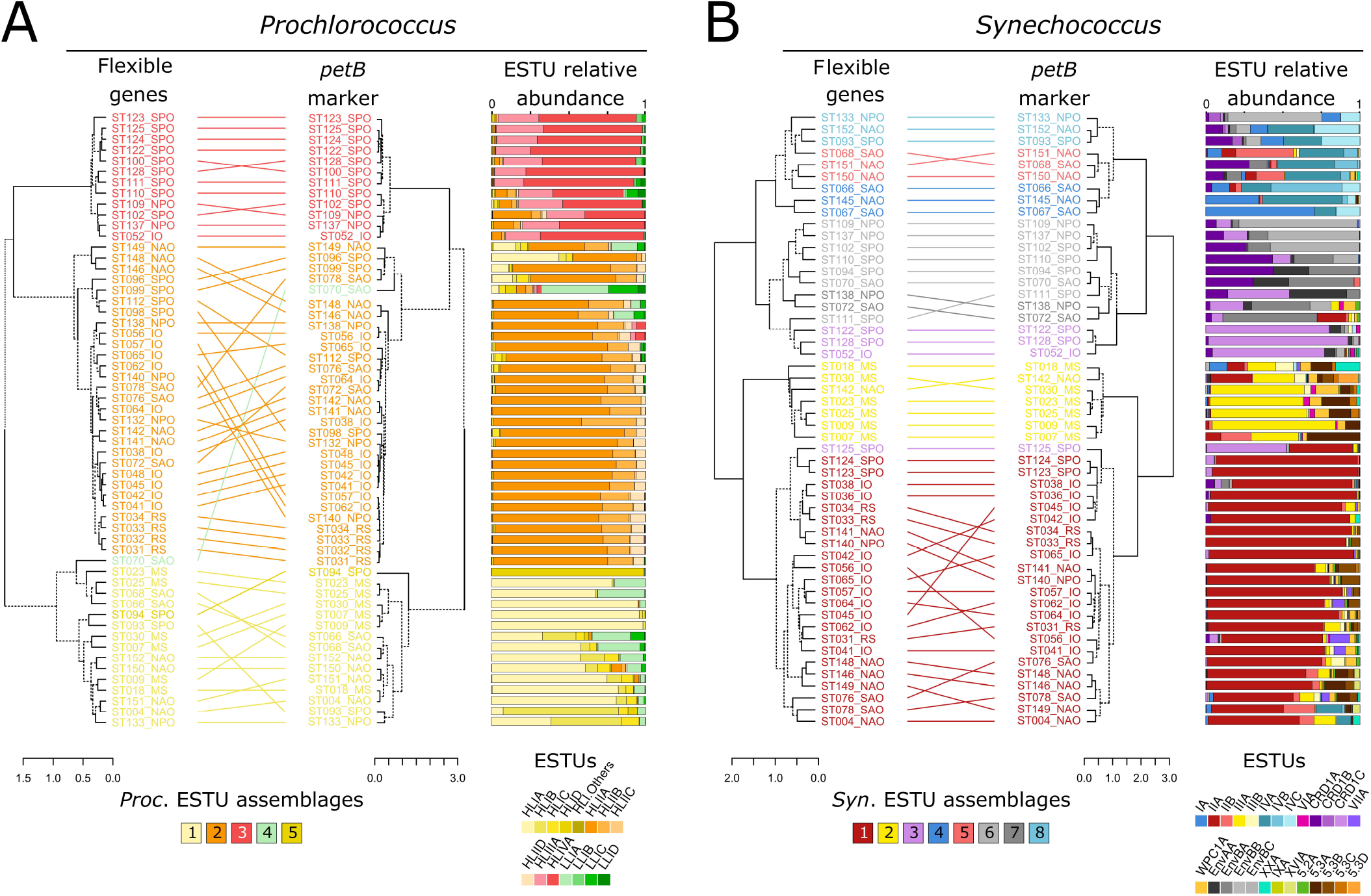
Comparison of clustering based on relative abundance profiles of ecologically significant taxonomic units (ESTUs) and of flexible genes for both picocyanobacteria. A. *Prochlorococcus*. B. *Synechococcus*. Leaves of the trees correspond to stations along the *Tara* Oceans transect that are colored according to the code shown at the bottom of the trees, corresponding to ESTU assemblages as determined by Farrant et al. (2016) by clustering stations exhibiting similar ESTU relative abundance profiles shown here on the right of each tree. ESTUs were colored according to the palette below each panel. Dotted lines in dendrograms indicate discrepancies between tree topologies. Accessory genes correspond to all genes except those defined as large-core genes in a previous study (9). Of note, due to a slightly different clustering method (cf. materials and methods), assemblage 7 (dark grey stations in 1B), which was discriminated from assemblage 6 in the Farrant et al. (2016) now clusters with this assemblage. Abbreviations: IO, Indian Ocean; MS, Mediterranean Sea; NAO, North Atlantic Ocean; NPO, North Pacific Ocean; RS, Red Sea; SAO, South Atlantic Ocean; SO, Southern Ocean.

### Distribution of flexible genes is tightly linked to environmental parameters and ESTUs

In order to reduce the amount of data and better interpret the global distribution of picocyanobacterial gene content, a correlation network of genes was built for each genus based on relative abundance profiles of genes across *Tara* Oceans samples. Its analysis emphasized four main modules of genes for *Prochlorococcus* (Fig. S1A) and five main modules for *Synechococcus* (Fig. S1B), each gene module being abundant in a different set of stations. These modules were then associated with the available environmental parameters (Figs. 2A-B) and to the relative abundance of *Prochlorococcus* or *Synechococcus* ESTUs at each station (Figs. 2C-D). For instance, the *Prochlorococcus brown* module was strongly correlated with nutrient concentrations, particularly nitrate and phosphate, and strongly anti-correlated with Fe availability (Fig. 2A). This module thus corresponds to genes preferentially found in Fe-limited high-nutrient low-chlorophyll (HNLC) areas. Indeed, the *brown* module *eigengenes* (Fig. S1A), representative of the abundance profiles of genes of this module at the different *Tara* Oceans stations, showed the highest abundances at stations *TARA*-100 to 125, localized in the South and North Pacific Ocean, as well as at *TARA*-052, a station located close to the northern coast of Madagascar and likely influenced by the Indonesian throughflow originating from the tropical Pacific Ocean (22, 31). Furthermore, the correlation of the *Prochlorococcus brown* module with the relative abundance of ESTUs at each station showed that it is also strongly associated with the presence of HLIIIA and HLIVA (Fig. 2C), previously shown to constitute the dominant *Prochlorococcus* ESTUs in low-Fe environments (22, 32, 33) but also the LLIB ESTU, found to dominate the LLI population in these low-Fe areas (22). Altogether, this example and analyses of all other *Prochlorococcus* and *Synechococcus* modules (SI Text1) show that the communities colonizing cold, Fe-, N- and/or P-depleted niches possess specific gene repertoires potentially involved in their adaptation to these peculiar environmental conditions.

**Figure 2.**
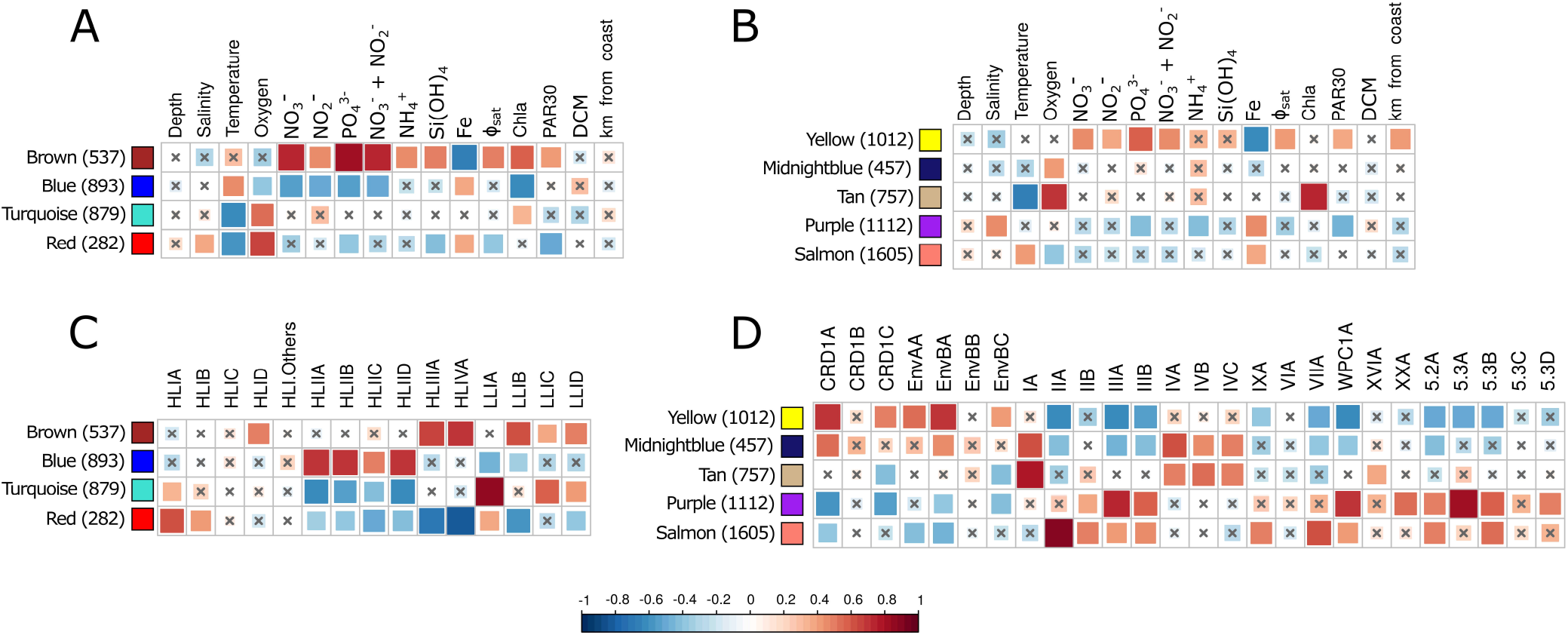
Correlation of picocyanobacterial module *eigengenes* to physico-chemical parameters and ESTU abundance. A, B. Correlation of module *eigengenes* to physico-chemical parameters for *Prochlorococcus* (A) and *Synechococcus* (B). C, D. Correlation of module *eigengenes* to relative abundance profiles of ESTUs *sensu* (Farrant et al., 2016). Pearson (A, B) and Spearman (B, D) correlation coefficient (R²) is indicated by the color scale. Each module is identified by a specific color and the number between brackets specifies the number of genes in each module. The *eigengene* is representative of the relative abundance of genes of a given module at each *Tara* Oceans station. Non-significant correlations (Student asymptotic p-value > 0.01) are marked by a cross. Φsat: index of iron limitation derived from satellite data. PAR30: satellite-derived photosynthetically available radiation at the surface, averaged on 30 days. DCM: depth of the deep chlorophyll maximum.

### Identification of individual genes potentially involved in niche partitioning

In order to identify flexible genes related to particular environmental conditions and to specific ESTU assemblages, we correlated relative abundance profiles of each gene to the eigengene vector of its corresponding module in order to identify the most representative genes of each module and thus the genes specifically present (or absent) in a given set of stations (Dataset 5, Figs. 3 and S2). Most genes retrieved this way encode proteins of unknown or hypothetical function (85.7% of 7,485 genes). Still, among the genes with a functional annotation (Dataset 6), a large fraction seems to have a function related to their realized environmental niche (Figs. 3 and S2). For instance, many genes involved in the transport and assimilation of nitrite and nitrate (*nirA, nirX, moaA-C, moaE, mobA, moeA, narB, M, nrtP*; all part of the same genomic island: Pro_GI004; (9)) as well as cyanate, an organic form of nitrogen (*cynA, B, D, S;* part of Pro_GI033), are enriched in the *Prochlorococcus* blue module, which is correlated with the HLIIA-D ESTU and to low inorganic N, P and Si levels and anti-correlated with Fe availability (Fig. 2A-C). This is consistent with previous studies showing that while few *Prochlorococcus* strains in culture possess the *nirA* gene and even less the *narB* gene, natural *Prochlorococcus* populations inhabiting N-poor areas do possess one or both of these genes (34–36). Similarly, numerous genes among the most representative genes of *Prochlorococcus brown*, red and *turquoise* modules are related to adaptation of HLIIIA/IVA, HLIA and LLIA ESTUs to Fe-limited, cold P-limited and cold, mixed waters, respectively (Fig. 3), and comparable results were obtained for *Synechococcus*, although the niche delineation was fuzzier than for *Prochlorococcus* at the module level (Fig. S2). These results therefore constitute a proof of concept that this network analysis was able to retrieve niche-related genes from metagenomics data.

**Figure 3.**
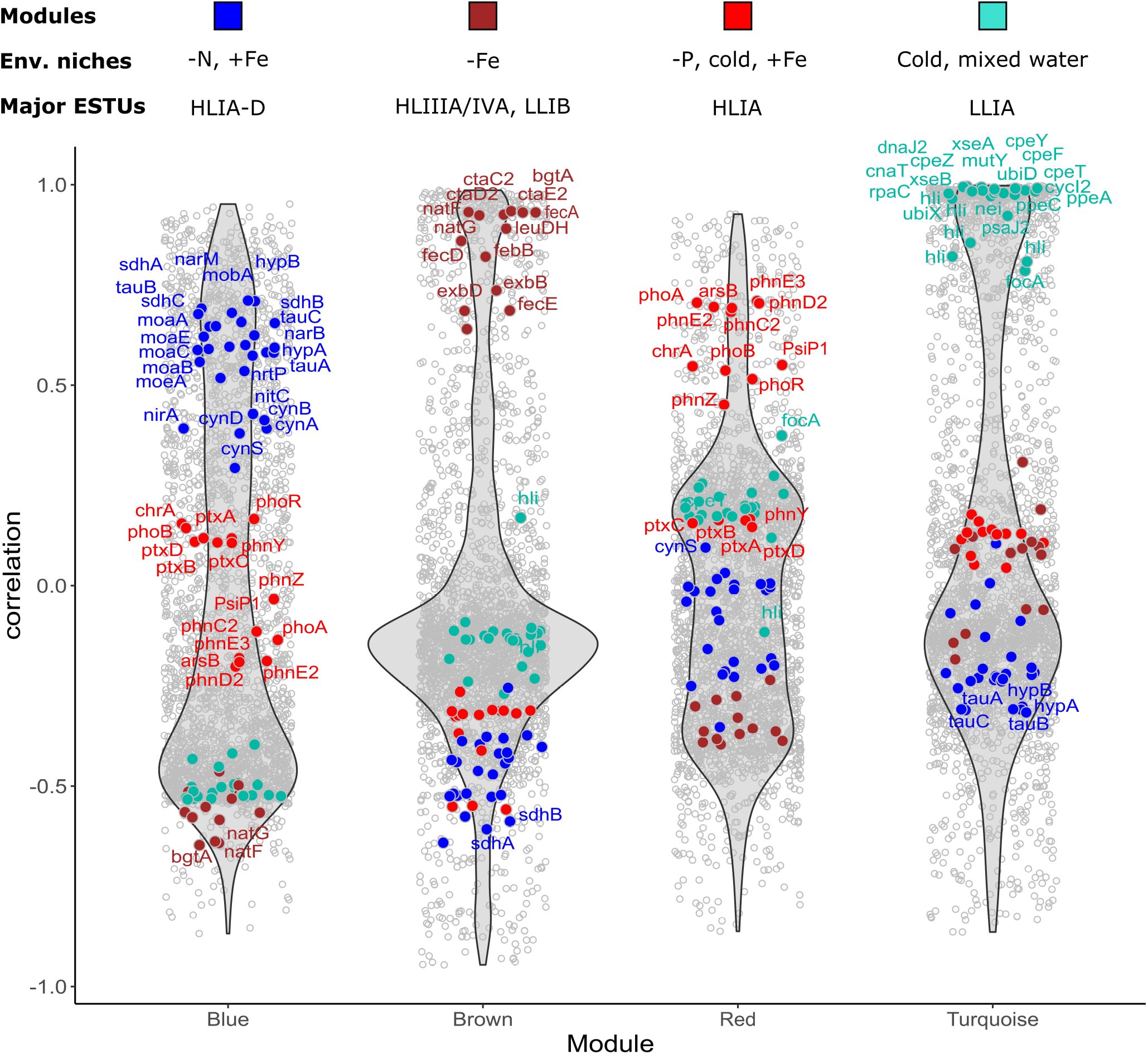
Violin plots highlighting the most representative genes of each *Prochlorococcus* module. For each module, each gene is represented as a dot positioned according to its correlation with the eigengene for each module, the most representative genes being localized on top of each violin plot. Genes mentioned in the text and/or in Dataset 6 have been colored according to the color of the corresponding module, indicated by a colored bar above each module. The text above violin plots indicates the most significant environmental parameter(s) and/or ESTU(s) for each module, as derived from Fig. 2.

### Identification of CAGs potentially involved in niche partitioning

In order to better understand the function of niche-related genes, notably the numerous ones encoding conserved hypothetical proteins, we then integrated these data with knowledge on the gene synteny in reference genomes using a network approach (Datasets 7 and 8). This led us to identify clusters of adjacent genes in reference genomes, several not previously reported in the literature, encompassing genes with similar distribution and abundance *in situ* and thus potentially involved in the same metabolic pathway (Figs. 4, S3 and S4; Dataset 6). Hereafter, these ecologically representative clusters of adjacent genes will be called ‘CAGs’.

**Figure 4.**
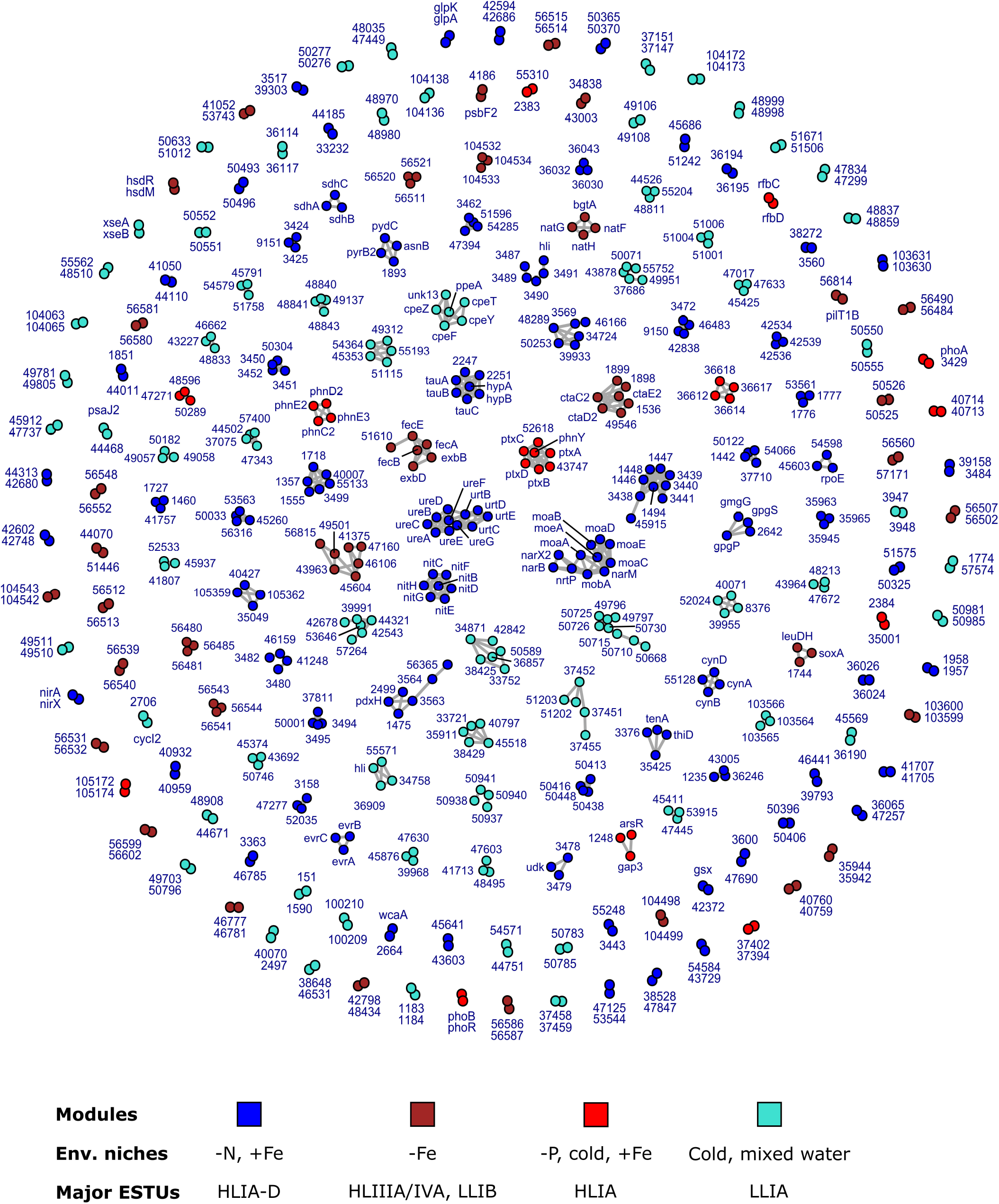
Delineation of *Prochlorococcus* CAGs, defined as a set of genes that are both adjacent in reference genomes and share a similar *in situ* distribution. Nodes correspond to individual genes with their gene name (or significant numbers of the CK number, e.g. 1234 for CK_00001234) and are colored according to their WGCNA module. A link between two nodes indicates that these two genes are less than 5 genes apart in at least one genome. The bottom insert shows the most significant environmental parameter(s) and/or ESTU(s) for each module, as derived from Fig. 2.

Regarding nitrogen, the well-known nitrate/nitrite gene cluster involved in uptake and assimilation of inorganic forms of nitrogen (see above) is present in most *Synechococcus* genomes (Dataset 6) and expectedly not restricted to a particular niche in natural *Synechococcus* populations, as shown by its quasi-absence from Weighted Correlation Network Analysis (WGCNA) modules. In *Prochlorococcus*, this cluster is separated into two CAGs, most genes being included in ProCAG_002, present in only 13 out of 118 *Prochlorococcus* genomes, while *nirA* and *nirX* form an independent CAG (ProCAG_001) due to their presence in many more genomes. Both CAGs are particularly enriched in *Prochlorococcus* populations thriving in low-N areas (Fig. S5A-B), as previously demonstrated by several authors (34–36). In *Prochlorococcus*, the quasi-core *ureA-G/urtB-E* genomic region was also found as a CAG (ProCAG_003) since it was comparatively impoverished in low-Fe compared to other regions (Fig. S5C-D) in agreement with its presence in only two out of six HLIII/IV genomes. In addition, we also uncovered several other *Prochlorococcus* and *Synechococcus* CAGs that seem to be involved in the transport and/or assimilation of more unusual and/or complex forms of nitrogen, including guanidine, cyanate, cyanide and possibly pyrimidine, which might either be degraded into elementary N, P or Fe molecules or possibly directly used by the cells for e.g. the biosynthesis of proteins or DNA. Indeed, we detected in both genera a CAG (ProCAG_004 and SynCAG_001; Figs. S6A-B, Dataset 6) that encompasses *speB2*, an ortholog of *Synechocystis* PCC 6803 *sll1077*, previously annotated as encoding an agmatinase (23, 37) and which was recently characterized as a guanidinase that degrades guanidine rather than agmatine to urea and ammonium (38). Interestingly *E. coli*, and likely other microorganisms as well, produce guanidine under nutrient-poor conditions, suggesting that guanidine metabolism is biologically significant and prevalent in natural environments (38, 39). Furthermore, the *ykkC* riboswitch candidate, which was shown to specifically sense guanidine and to control the expression of a variety of genes involved in either guanidine metabolism or nitrate, sulfate, or bicarbonate transport, is located immediately upstream of this CAG in *Synechococcus* reference genomes, all genes of this cluster being predicted by RegPrecise 3.0 to be regulated by this riboswitch (Fig. S6C; (39, 40)). The presence of *hypA* and B homologs within this CAG furthermore suggests that, in the presence of guanidine, the latter could be involved in the insertion of Ni_2_^+^, or another metal cofactor, in the active site of guanidinase. Additionally, we speculate that the next three genes encoding an ABC transporter, similar to the TauABC taurine transporter in *E. coli* (Fig. S6C), could be involved in guanidine transport in low-N areas. Of note, the presence of a gene encoding a putative Rieske Fe-sulfur protein (CK_00002251), downstream of this gene cluster in most *Synechococcus*/*Cyanobium* genomes possessing this CAG, seems to constitute a specificity compared to its homologs in *Synechocystis* sp. PCC 6803 and might explain why this CAG is absent from picocyanobacteria thriving in low-Fe areas, while it is present in a large proportion of the population in most other oceanic areas (Figs. S6A-B).

As concerns compounds containing a cyano radical (C⍰N), the cyanate transporter genes (*cynABD*) are scarce in both *Prochlorococcus* (present only in two HLI and five HLII genomes) and *Synechococcus* genomes (mostly in clade III strains; (9, 41, 42)). In the field, a small proportion of the *Prochlorococcus* community possesses the corresponding CAG (ProCAG_005; Fig. S7A-B), also including the conserved hypothetical gene CK_00055128, in warm, Fe-replete waters, while these genes were not included in a module, and thus not in a CAG, in *Synechococcus* (Dataset 6; Fig. S7C). Interestingly, we also uncovered a 7-gene CAG (ProCAG_006 and SynCAG_002), encompassing a putative nitrilase gene (*nitC*), which also suggests that most *Synechococcus* cells and a more variable proportion of the *Prochlorococcus* population could use nitriles or cyanides in warm, Fe-replete waters and more particularly in low-N areas such as the Indian Ocean (Fig. 5A-B). The whole operon *(nitHBCDEFG;* Fig. 5C), called Nit1C, was shown to be upregulated in the presence of cyanide and to trigger an increase in the rate of ammonia accumulation in the heterotrophic bacterium *Pseudomonas fluorescens* (43), suggesting that like cyanate, cyanide could constitute an alternative nitrogen source in marine picocyanobacteria as well. Yet, given the potential toxicity of these C⍰N-containing compounds, we cannot exclude that these CAGs could also be devoted to cell detoxification (39, 41), as it is the case for arsenate and chromate (44, 45), which act as analogs of phosphate and sulfate respectively, and are toxic to marine phytoplankton (46).

**Figure 5.**
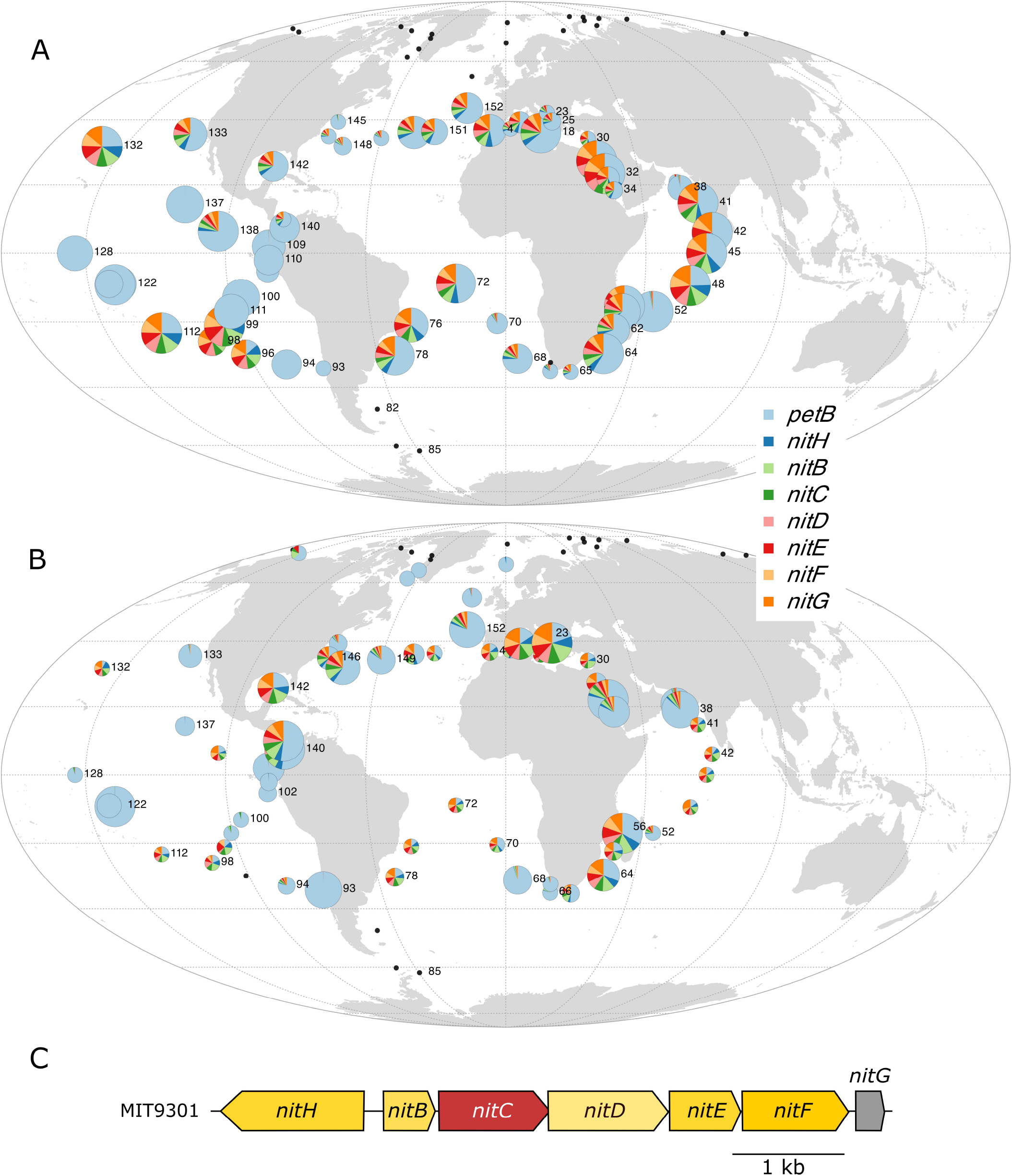
Global distribution map of CAG involved in nitriles or cyanides transport and assimilation. (*A*) *Prochlorococcus* (ProCAG_006) and (*B*) *Synechococcus* SynCAG_002. (*C*) Genomic region in *Prochlorococcus marinus* MIT9301. The size of the circle is proportional to relative abundance of *Prochlorococcus* as estimated based on the single-copy core gene *petB* gene and this gene was also used to estimate the relative abundance of other genes in the population.

Also noteworthy is the presence of a CAG encompassing *asnB, pyrB2* and *pydC* (ProCAG_007, SynCAG_003, Fig. S8), which could contribute to an alternative pyrimidine biosynthesis pathway and thus provide another way for cells to recycle complex nitrogen forms. While this CAG is found in only one fifth of HLII genomes and in quite specific locations for *Prochlorococcus*, notably in the Red Sea, it is found in most *Synechococcus* cells in warm, Fe-replete, N and P-depleted niches, consistent with its phyletic pattern showing its absence only from most clade I, IV, CRD1 and EnvB genomes (Fig. S8; Dataset 6). More generally, most N-uptake and assimilation genes in both genera were specifically absent from Fe-depleted areas, including the *nirA*/*narB* CAG for *Prochlorococcus*, as mentioned by Kent et al. (30) as well as guanidinase and nitrilase CAGs. In contrast, picocyanobacterial populations present in low-Fe areas possess, in addition to the core ammonium transporter *amt1*, a second transporter *amt2*, also present in cold areas for *Synechococcus* (Fig. S9). Additionally, *Prochlorococcus* populations thriving in HNLC areas also possess two amino acid-related CAGs that are quasi-core in *Synechococcus*, the first one involved in polar amino acids N-II transport system (ProCAG_008; *natF-G-H-bgtA;* (47); Fig. S10A-B) and the second one (*leuDH, soxA*, CK_00001744, ProCAG_009, Fig. S10C-D) that notably encompasses a leucine dehydrogenase, able to produce ammonium from branched-chain amino acids. Thus, the primary nitrogen sources for picocyanobacterial populations dwelling in Fe-limited areas seem to be ammonium and amino acids.

Adaptation to phosphorus depletion has been well documented in marine picocyanobacteria showing that while in P-replete waters *Prochlorococcus* and *Synechococcus* essentially rely on inorganic phosphate acquired by core transporters (PstABC), strains isolated from low-P regions and natural populations thriving in these areas additionally contain a number of accessory genes related to P metabolism, located in specific genomic islands (9, 25– 28, 48). Here, we indeed found that virtually the whole *Prochlorococcus* population in the Mediterranean Sea, the Gulf of Mexico and the Western North Atlantic Ocean, which are known to be P-limited (26, 49), contained the phoBR operon (ProCAG_010, Fig. S11A-B) that encodes a two-component system response regulator, as well as the ProCAG_011, including the alkaline phosphatase *phoA*. By comparison, in *Synechococcus*, we only identified the *phoBR* CAG (SynCAG_005, Fig. S11C) that is systematically present in warm waters whatever their limiting nutrient, in agreement with its phyletic pattern in reference genomes showing its specific absence from cold thermotypes (clades I and IV, Dataset 6). Furthermore, although our analysis did not retrieve them within CAGs due to the variability of the content and order of genes in this genomic region, even within each genus, several other P-related genes were enriched in low-P areas but interestingly partially differed between *Prochlorococcus* and *Synechococcus* (Figs. S11, 3, S2 and Dataset 6). While the genes putatively encoding a chromate transporter (ChrA) and an arsenate efflux pump ArsB were present in both genera in different proportions, a putative transcriptional phosphate regulator related to PtrA (CK_00056804; (50)) was specific to *Prochlorococcus. Synechococcus* in contrast harbors a large variety of alkaline phosphatases (PhoX, CK_00005263 and CK_00040198) as well as the phosphate transporter SphX (Fig. S11).

A second alternative P form are phosphonates, i.e. reduced organophosphorus compounds containing C–P bonds, which constitute up to 25% of the high-molecular-weight dissolved organic P pool in the open ocean (51). Indeed, the quasi-totality of the *Prochlorococcus* population of the most P-limited areas of the ocean possess, additionally to the core phosphonate ABC transporter (*phnD1-C1-E1*), a second previously unreported putative phosphonate transporter (*phnC2-D2-E2-E3*; ProCAG_012; Fig. 6A), while these genes are only present in a few *Prochlorococcus* (including MIT9314) and no *Synechococcus* genomes. Furthermore, as previously mentioned in several studies (52–54), a fairly low proportion of *Prochlorococcus* populations thriving in low-P areas also possess a gene cluster encompassing the *phnYZ* operon, involved in C-P bond cleavage, the putative phosphite dehydrogenase *ptxD* as well as the phosphite and methylphosphonate transporter *ptxABC* (ProCAG_0013, Dataset 6, and Fig. 6B, (54–56)). Compared to these previous studies that mainly reported the presence of these genes in *Prochlorococcus* cells from the North Atlantic Ocean, here we show that they actually occur in a much larger geographic area, including the Mediterranean Sea, the Gulf of Mexico and the ALOHA station (*TARA*_132) in the North Pacific, and are also present in an even larger proportion of the *Synechococcus* population (Fig. S12, Dataset 6). Interestingly, *Synechococcus* cells from the Mediterranean Sea, dominated by clade III, seem to lack *phnYZ*, in agreement with the phyletic pattern of these genes in reference genomes, showing the absence of this two-gene operon in the sole clade III strain that possesses the *ptxABDC* gene cluster. In contrast, the presence of the complete gene set (*ptxABDC*-*phnYZ*) in the North Atlantic and at the entrance of the Mediterranean Sea as well as in several clade II reference genomes rather suggests that it is primarily attributable to this specific clade. Altogether, our data indicate that at least part of the natural populations of both *Prochlorococcus* and *Synechococcus* would be able to assimilate phosphonate and phosphite as alternative P-sources in low-P areas using the ptxABDC-phnYZ operon. Yet, the fact that no picocyanobacterial genome except *P. marinus* RS01 (Fig. 6C) possesses both *phnC2-D2-E2-E3* and *phnYZ*, raises the question of how the phosphonate taken up by the *phnC2-D2-E2-E3* transporter is metabolized in these cells. Finally, although the Mediterranean Sea is not known to be N-limited, all reference clade III genomes possess a complete set of genes involved in the assimilation of organic nitrogen (Dataset 6), suggesting that at least part of these organic nutrients might also be a source of organic phosphorus.

**Figure 6.**
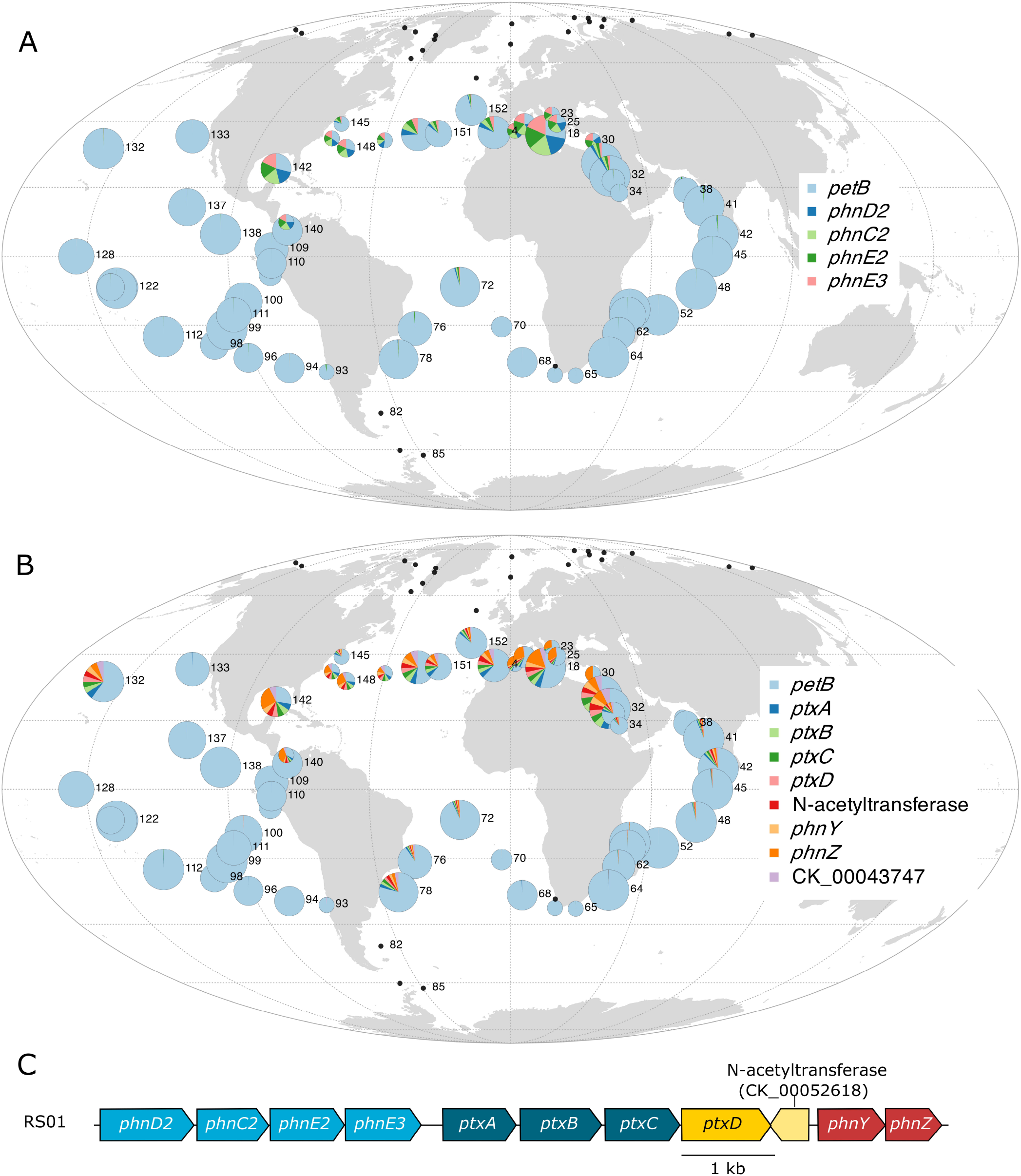
Global distribution map of CAGs putatively involved in phosphonate and phosphite transport and assimilation. *Prochlorococcus* (*A*) ProCAG_012 putatively involved in phosphonate transport, (*B*) ProCAG_013, involved in phosphonate/phosphite uptake and assimilation and phosphonate C-P bond cleavage, (*C*) The genomic region encompassing both *phnC2-D2-E2-E3* and *ptxABDC-phnYZ* specific to *P. marinus* RS01. The size of the circle is proportional to relative abundance of *Prochlorococcus* as estimated based on the single-copy core gene *petB* and this gene was also used to estimate the relative abundance of other genes in the population.

As for macronutrients, it has been hypothesized that the survival of marine picocyanobacteria in low-Fe regions was made possible through several strategies, including the elimination from the genomes of genes encoding proteins that contain Fe as a cofactor, the replacement of Fe by another metal cofactor, and the acquisition of genes involved in Fe uptake and storage (24, 25, 30, 33, 57). Accordingly, several CAGs encompassing genes encoding proteins interacting with Fe were found in the present study to be anti-correlated with HNLC regions in both genera. These include three subunits of the (photo)respiratory complex succinate deshydrogenase (SdhABC, ProCAG_014, SynCAG_006, Fig. S13; (58)) as well as Fe-containing proteins encoded in most of the abovementioned CAGs involved in N or P metabolism, such as the guanidinase CAG (Fig. S6), the NitC1 CAG (Fig. 5), the *pyrB2* CAG (Fig. S8), the phosphonate CAGs (Figs. 6 and S12) and the urea and inorganic nitrogen CAGs (Fig. S5). Most *Synechococcus* cells thriving in Fe-replete areas also possess the *sodT/sodX* CAG (SynCAG_007, Fig. S14A-B) involved in nickel transport and maturation of the Ni-superoxide dismutase (SodN), these three genes being in contrast core in *Prochlorococcus*. Additionally, *Synechococcus* from Fe-replete areas, notably from the Mediterranean Sea and the Indian Ocean, specifically possess two CAGs (Syn CAG_008 and 009; Fig. S14C-D), involved in the biosynthesis of a polysaccharide capsule that appear to be most similar to the *E. coli* groups 2 and 3 *kps* loci (59). These extracellular structures, known to provide protection against biotic or abiotic stress, were recently shown in *Klebsiella* to provide a clear fitness advantage in nutrient-poor conditions since they were associated with increased growth rates and population yields (60). Yet, while these authors suggested that capsules may play a role in Fe uptake, the significant reduction of the relative abundance of *kps* genes in low-Fe compared to Fe-replete areas (t-test p-value <0.05 for all genes of the Syn CAG_008 and 009 except CK_00002157; Fig. S14C) and their absence in CRD1 strains (Dataset 6) rather suggests that these capsules may be too energy-consuming for some picocyanobacteria thriving in this peculiar niche, while they may have a meaningful and previously overlooked role in their adaptation to low-P and low-N niches.

A number of CAGs were in contrast found to be enriched in populations dwelling in HNLC environments, dominated by *Prochlorococcus* HLIIIA/HLIVA/LLIB and *Synechococcus* CRD1A/EnvBA ESTUs (Fig. 2). For *Prochlorococcus*, this includes the abovementioned *natFGH* (ProCAG_008) and *leudH/soxA* (ProCAG_009) CAGs, involved in amino acid metabolism (Fig. S10), while a large proportion of the *Synechococcus* populations in these areas possess i) a large CAG involved in glycine betaine synthesis and transport (SynCAG_010, Fig. S15A-B; (9, 61)), almost absent from low-N areas, ii) a CAG encompassing a flavodoxin and a thioredoxin reductase (SynCAG_011, Fig. S15C-D), mostly absent from low-P areas, as well as iii) the *nfeD*-*floT1-floT2 CAG* (SynCAG_012, Fig. S16A-B) involved in the production of lipid rafts, potentially affecting cell shape and motility (9, 62). Both *Prochlorococcus* and *Synechococcus* thriving in low-Fe waters also possess the TonB-dependent siderophore uptake operon (*fecDCAB-tonB-exbBD*, Dataset 6). The latter gene cluster, which is found in a few picocyanobacterial genomes and was previously shown to be anti-correlated with dissolved Fe concentration (24, 25, 57), is indeed systematically present in a significant part of the *Prochlorococcus* and *Synechococcus* population in low-Fe areas (ProCAG_015 and SynCAG_013-014, Fig. S17). However, it is also present in a small fraction of the populations thriving in the Indian Ocean, consistent with its occurrence in two *Prochlorococcus* HLII and one *Synechococcus* clade II genomes (Dataset 6). The most striking result in this category is that the vast majority of *Prochlorococcus* cells thriving in low-Fe regions possess a CAG encompassing the *ctaC2-D2-E2* operon, also found in 85% of all *Synechococcus* reference genomes, including all CRD1 (Fig. 7; Dataset 6). This CAG encodes the alternative respiratory terminal oxidase ARTO, a protein complex that has been suggested to be part of a minimal respiratory chain in the cytoplasmic membrane, potentially providing an additional electron sink under Fe-deprived conditions (63, 64). Furthermore, a *Synechocystis* mutant in which the *ctaD2* and *ctaE2* genes had been inactivated was found to display markedly impaired Fe reduction and uptake rates as compared to wild-type cells, suggesting that ARTO is involved in the reduction of Fe(III) into Fe(II) prior to its transport through the plasma membrane via the Fe(II) transporter FeoB (65). Thus, the presence of the ARTO system appears to represent a major and previously unreported adaptation for *Prochlorococcus* populations thriving in low-Fe areas.

**Figure 7.**
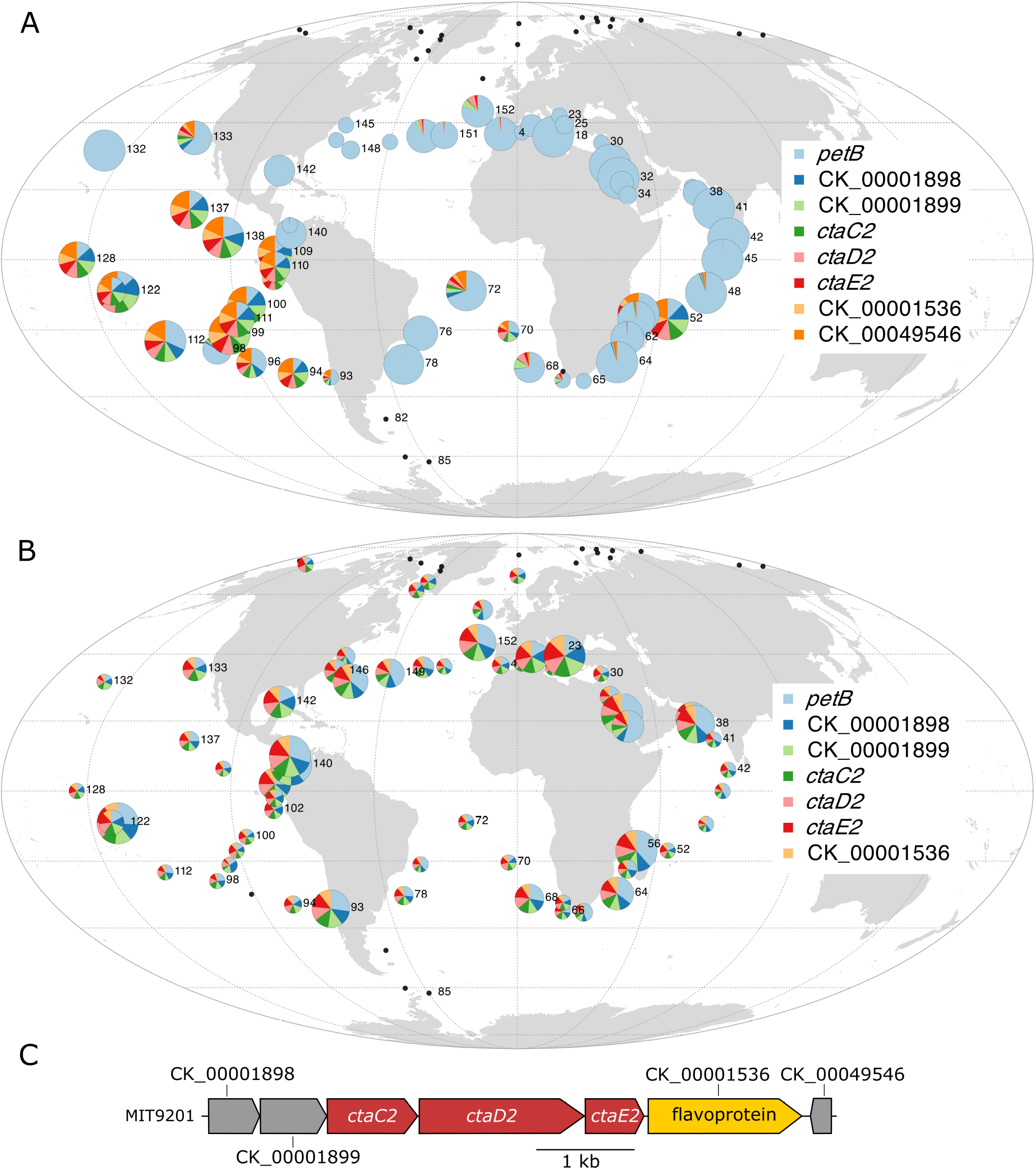
Global distribution map of the *Prochlorococcus* CAGs involved in the biosynthesis of an alternative respiratory terminal oxidase (ARTO). (*A*) *Prochlorococcus* ProCAG_016, (*B*) *Synechococcus* SynCAG_015. The size of the circle is proportional to relative abundance of *Prochlorococcus* as estimated based on the single-copy core gene *petB* and this gene was also used to estimate the relative abundance of other genes in the population.

Besides genes involved in nutrient acquisition and metabolism, several *Prochlorococcus* and *Synechococcus* CAGs were found to be correlated with low-temperature waters. A closer examination of *Prochlorococcus* CAGs however, shows that their occurrence is not directly related to temperature adaptation but mainly explained by the prevalence at high latitude of either i) the HLIA ESTU (Fig. 2A, C and Fig. 4), the red module encompassing most of the abovementioned CAGs involved in P-uptake and assimilation pathways, or ii) the LLIA ESTU, present in surface waters at vertically-mixed stations, the *turquoise* module mainly gathering *Prochlorococcus* LL-specific genes, such as Pro_CAG_017, involved in phycoerythrin-III biosynthesis (*ppeA, cpeFTZY, unk13*) or ProCAG_018, encoding the two subunits of exodeoxyribonuclease VII (XseA-B). As concerns *Synechococcus*, although a fairly high number of CAGs were identified in the tan module associated with ESTUs IA and IVA-C (Fig. 2B, D and Fig. S4), only very few are conserved in more than two reference strains and/or have a characterized function (Dataset 6). Among these, at least one CAG is clearly related to adaptation to cold waters, the orange carotenoid protein (OCP) operon (*ocp-crtW-frp*; SynCAG_016). Indeed, this operon is involved in a photoprotective process (66) and was recently shown to provide cells with the ability to deal with oxidative stress under cold temperatures (67). In agreement with the latter study, our data shows that *Synechococcus* populations colonizing mixed waters at high latitudes or in upwelling areas all possess the ocp CAG (Fig. S18), highlighting the importance of this photoprotection system in *Synechococcus* populations colonizing cold and temperate areas. *Synechococcus* populations thriving in cold waters also appear to be enriched in CAGs involved in gene regulation. This includes transcriptional regulators involved in the regulation of the CA4-A form of the type IV chromatic acclimation process (*fciA-B;* SynCAG_017), consistent with the predominance of *Synechococcus* CA4-A cells in temperate or cold environments (68–70)(Dataset 6) as well as the *hidABC* operon (SynCAG_018), involved in the synthesis of a secondary metabolite (hierridin C; (71)). Altogether, the fairly low number of ‘strong’ CAGs associated with temperature supports the hypothesis that adaptation to cold temperature is not mediated by evolution of gene content but rather of protein sequences (8, 9, 30, 72).

In conclusion, our analysis of *Prochlorococcus* and *Synechococcus* gene distributions at the global scale using the deeply sequenced metagenomes collected along the *Tara* Oceans expedition transect revealed that each community has a specific gene repertoire, with different sets of accessory genes being highly correlated with distinct ESTUs and physicochemical parameters. As previously suggested for *Prochlorococcus* (30), this strong correlation between taxonomy and gene content strengthens the idea that, in both genera, genome evolution mainly occurs by vertical transmission and selective gene retention, with a fairly low extent of lateral gene transfer between clades. By combining information about gene synteny in 256 reference genomes with the distribution and abundance of these genes in the field, we further managed to delineate suites of adjacent genes likely involved in the same metabolic pathways that may have a crucial role in adaptation to specific niches. These analyses confirmed previous observations about the niche partitioning of individual genes and a few genomic regions involved in nutrient uptake and assimilation (24, 25, 27, 30, 34, 36). Most importantly, this network approach unveiled several novel genomic regions that could confer cells a fitness benefit in particular niches and also highlighted that some previously detected individual genes are part of larger genomic regions. It notably revealed the potential importance of the uptake and assimilation of organic forms of limiting nutrients, which might either be directly used by the cells, e.g. for the biosynthesis of proteins or DNA, or be degraded into inorganic N and/or P forms. Consistently, many CAGs potentially involved in the uptake and assimilation of complex compounds, such as guanidine, C⍰N-containing compounds or pyrimidine were present in both N- and P-depleted waters, and might constitute an advantage in areas of the world ocean co-limited in these nutrients (26). In contrast, most of these CAGs were specifically absent from N and/or P-rich, Fe-poor areas ((30); this study). Adaptation to Fe-limitation seemingly relies on specific adaptation mechanisms including reduction of Fe^3+^ to Fe^2+^ using ARTO, Fe storage, Fe scavenging using siderophores as well as reduction of the iron quota and of energy-consuming adaptation mechanisms, such as polysaccharide capsule biosynthesis. Altogether, this study provides unique insights into the functional basis of microbial niche partitioning and the molecular bases of fitness in key members of the phytoplankton community. A future challenge will clearly consist of biochemically characterizing the function of the different genes, including many unknown, gathered in the above-mentioned CAGs (Datasets 5 and 6), which are sometimes present only in a few or even a single strain but can occur in a large part or even the whole *Prochlorococcus* and/or *Synechococcus* population *in situ*, and which likely all contribute to the same complex and/or metabolic pathway.

## Materials and Methods

### *Tara* Oceans dataset

A total of 131 bacterial-size metagenomes (0.2-1.6 µm for stations *TARA*_004 to *TARA*_052 and 0.2-3µm for *TARA*_056 to *TARA*_152), collected in surface from 83 stations along the *Tara* Oceans expedition transect (73), were used in this study. Briefly, all metagenomes were sequenced as Illumina overlapping paired reads of 100-108 bp and paired reads were merged and trimmed based on quality, resulting in 100-215 bp fragments, as previously described (22). All metagenomes and corresponding environmental parameters were retrieved from PANGAEA (www.pangaea.de/) except for Fe and ammonium concentrations that were modeled and the Fe limitation index Φ_sat_ that was calculated from satellite data, as previously described (22).

### Recruitment and taxonomic and functional assignment of metagenomic reads

Metagenomic reads were first recruited against 256 reference genomes, including the 97 genomes available in the information system Cyanorak *v2.1* (www.sb-roscoff.fr/cyanorak; (28)) as well as 84 additional WGS, 27 MAGs and 48 SAGs retrieved from Genbank (Dataset 9). Recruitment was made using MMseqs2 Release 11-e1a1c (76) with maximum sensitivity (mmseqs search -s 7.5) and limiting the results to one target sequence (mmseqs filterdb -- extract-lines 1). Using the same MMseqs2 options, the resulting reads were then mapped to an extended database of 978 genomes, including all picocyanobacterial reference genomes complemented with 722 outgroup cyanobacterial genomes downloaded from NCBI. Reads mapping to outgroup sequences or having less than 90% of their sequence aligned were filtered out and the remaining reads were taxonomically assigned to either *Prochlorococcus* or *Synechococcus* according to their best hit. Reads were then functionally assigned to a cluster of likely orthologous genes (CLOGs) from the information system Cyanorak *v2.1* based on the position of their MMseqs2 match on the genome, the coordinates of which correspond to a particular gene in the database. More precisely, a read was functionally assigned to a gene if at least 75% of its size was aligned to the reading frame of this gene and if the percentage identity of the blast alignment was over 80%. Finally, read counts were aggregated by CLOG and normalized by dividing read counts by L-l+1, where L represents the average gene length of the CLOG and l the mean length of recruited reads. Only environmental samples that contained at least 2,500 and 1,700 distinct CLOGs for *Synechococcus* and *Prochlorococcus*, respectively, were kept, corresponding roughly to the average number of genes in a *Synechococcus* and a *Prochlorococcus* HL genome, respectively. After this filtration step, a CLOG was kept if it showed a gene-length normalized abundance higher than 1 (i.e., a gene coverage of 1) in at least 2 of the selected environmental samples. Then, large-core genes, as previously defined (9), were removed to keep only accessory genes. The resulting abundance profiles were used to perform co-occurrence analyses by weighted genes correlation network analysis, as detailed below (WGCNA, (74)).

### Station clustering and ESTU analyses

In order to cluster stations displaying similar CLOG abundance patterns, the abundance of a given CLOG in a sample was divided by the total CLOG abundance in this sample to obtain relative abundance profiles per sample. Bray-Curtis similarities were calculated from these profiles and used to cluster *Tara* Oceans stations with the Ward’s minimum variance method (75). The same normalization method was applied to picocyanobacterial ESTUs that were defined based on the *petB* marker gene at each station using a similar approach as in Farrant et al. (2016) but using a Ward’s minimum variance method (75) to be consistent with the clustering of CLOG profiles. In order to check whether the Bray-Curtis distances between stations based on petB picocyanobacterial communities and based on gene content were significantly correlated, a mantel test was performed between the distance matrices, as implemented in the R package *vegan* v2.5 with 9,999 permutations (76).

### Gene co-occurrence network analysis

A data-reduction approach based on WGCNA, as implemented in the R package WGCNA v1.51 (77), was used to build a co-occurrence network of CLOGs based on their relative abundance in *Tara* Oceans stations and to delineate modules of CLOGs (i.e., subnetworks). The WGCNA adjacency matrix was calculated in ‘signed’ mode (i.e., considering correlated and anti-correlated CLOGs separately), by using the *Pearson* correlation between pairs of CLOGs (based on their relative abundance in every sample) and raising it to the power 12, which allowed to obtain a scale-free topology of the network. Modules were identified by setting the minimum number of genes in each module to 100 and 50 for *Synechococcus* and *Prochlorococcus*, respectively, and by forcing every gene to be included in a module. The *eigengene* of each module (representative of the relative abundance of genes of a given module at each *Tara* Oceans station) was then correlated to environmental parameters and to the relative abundance of ESTUs. Finally, genes in each module with the highest correlation to the *eigengene* (a measurement called ‘membership’), were extracted in order to identify the most representative genes of each module.

### Identification of differentially distributed clusters of adjacent genes (CAGs)

Results on individual niche-related genes identified by WGCNA were then integrated with knowledge on gene synteny in reference genomes (Datasets 7 and 8). For each WGCNA module, we defined CAGs by searching adjacent genes of the module in the 256 reference genomes. In order to be considered as belonging to the same CAG, two genes of the same module must be less than 6 genes apart in 80% of the genomes in which these two genes are present. This led us to identify clusters of adjacent genes in reference genomes, comprising genes displaying a similar distribution pattern, called CAGs. A network of CAGs was then built for each WGCNA module, taking into account the number of genomes in which these genes are adjacent (Figs. 4, S3 and S4). An unweighted, undirected graph was drawn for each module according to the Fruchterman-Reingold layout algorithm implemented in the R package igraph. This is a force-directed algorithm, meaning that node layout is determined by the forces pulling nodes together and pushing them apart. In other words, its purpose is to position the nodes of a graph so that the edges of more or less equal length are gathered together and to avoid as many crossing edges as possible.

## Data sharing plans

All genomic and metagenomic data used in this study are publicly available

## Acknowledgments

This work was supported by the French “Agence Nationale de la Recherche” Programs SAMOSA (ANR-13-ADAP-0010), CINNAMON (ANR-17-CE02-0014-01), EFFICACY (ANR-19-CE02-0019) and France Génomique (ANR-10-INBS-09) as well as the European Union program Assemble+ (Horizon 2020, under grant agreement number 287589). We acknowledge Christophe Six for his help with cloning some of the *Synechococcus* strains used in this study and Francisco M. Cornejo-Castillo for useful discussions. We also thank the support and commitment of the *Tara* Oceans coordinators and consortium, Agnès b. and E. Bourgois, the Veolia Environment Foundation, Région Bretagne, Lorient Agglomeration, World Courier, Illumina, the EDF Foundation, FRB, the Prince Albert II de Monaco Foundation, the *Tara* schooner, and its captains and crew. *Tara* Oceans would not exist without continuous support from 23 institutes (http://oceans.taraexpeditions.org).

## References

1. H. Tettelin, et al., Genome analysis of multiple pathogenic isolates of Streptococcus agalactiae: Implications for the microbial “pan-genome.” Proc Natl Acad Sci USA 102, 13950–13955 (2005).

2. M. López-Pérez, F. Rodriguez-Valera, Pangenome evolution in the marine bacterium Alteromonas. Genome Biol Evol 8, 1556–1570 (2016).

3. T. D. Read, et al., The genome sequence of Bacillus anthracis Ames and comparison to closely related bacteria. Nature 423, 81–86 (2003).

4. C. Zhu, T. O. Delmont, T. M. Vogel, Y. Bromberg, Functional basis of microorganism classification. PLOS Comput Biol 11, e1004472 (2015).

5. M. L. Reno, N. L. Held, C. J. Fields, P. V. Burke, R. J. Whitaker, Biogeography of the Sulfolobus islandicus pan-genome. Proc Natl Acad Sci USA 106, 8605–8610 (2009).

6. S. S. Porter, P. L. Chang, C. A. Conow, J. P. Dunham, M. L. Friesen, Association mapping reveals novel serpentine adaptation gene clusters in a population of symbiotic Mesorhizobium. ISME J 11, 248–262 (2017).

7. S. Kellner, et al., Genome size evolution in the Archaea. Emerg Top Life Sci 2, 595–605 (2018).

8. G. C. Kettler, et al., Patterns and implications of gene gain and loss in the evolution of Prochlorococcus. PLOS Genet 3, e231 (2007).

9. H. Doré, et al., Evolutionary mechanisms of long-term genome diversification associated with niche partitioning in marine picocyanobacteria. Front Microbiol 11 (2020).

10. N. Kashtan, et al., Single-cell genomics reveals hundreds of coexisting subpopulations in wild Prochlorococcus. Science 344, 416–420 (2014).

11. P. B. Pearman, A. Guisan, O. Broennimann, C. F. Randin, Niche dynamics in space and time. Trends Ecol Evol 23, 149–158 (2008).

12. T. O. Delmont, A. M. Eren, Linking pangenomes and metagenomes: the Prochlorococcus metapangenome. PeerJ 6, e4320 (2018).

13. J.-H. Hehemann, et al., Adaptive radiation by waves of gene transfer leads to fine-scale resource partitioning in marine microbes. Nat Commun 7, 12860 (2016).

14. H. Koch, et al., Genomic, metabolic and phenotypic variability shapes ecological differentiation and intraspecies interactions of Alteromonas macleodii. Sci Rep 10 (2020).

15. J. P. Engelberts, et al., Characterization of a sponge microbiome using an integrative genome-centric approach. ISME J 14, 1100–1110 (2020).

16. B. J. Tully, E. D. Graham, J. F. Heidelberg, The reconstruction of 2,631 draft metagenome-assembled genomes from the global oceans. Sci Data 5, 170203 (2018).

17. B. L. Hurwitz, A. H. Westveld, J. R. Brum, M. B. Sullivan, Modeling ecological drivers in marine viral communities using comparative metagenomics and network analyses. Proc Natl Acad Sci USA 111, 10714–10719 (2014).

18. A. Meziti, et al., Quantifying the changes in genetic diversity within sequence-discrete bacterial populations across a spatial and temporal riverine gradient. ISME J 13, 767–779 (2019).

19. I. Raimundo, et al., Functional metagenomics reveals differential chitin degradation and utilization features across free-living and host-associated marine microbiomes. Microbiome 9, 43 (2021).

20. P. Flombaum, et al., Present and future global distributions of the marine Cyanobacteria Prochlorococcus and Synechococcus. Proc Natl Acad Sci USA 110, 9824–9 (2013).

21. N. Visintini, A. C. Martiny, P. Flombaum, Prochlorococcus, Synechococcus, and picoeukaryotic phytoplankton abundances in the global ocean. Limnol Oceanogr 6, 207–215 (2021).

22. G. K. Farrant, et al., Delineating ecologically significant taxonomic units from global patterns of marine picocyanobacteria. Proc Natl Acad Sci USA 113, E3365–E3374 (2016).

23. A. G. Kent, et al., Parallel phylogeography of Prochlorococcus and Synechococcus. ISME J 13, 430–441 (2019).

24. N. A. Ahlgren, B. S. Belisle, M. D. Lee, Genomic mosaicism underlies the adaptation of marine Synechococcus ecotypes to distinct oceanic iron niches. Environ Microbiol 22, 1801– 1815 (2020).

25. C. A. Garcia, et al., Linking regional shifts in microbial genome adaptation with surface ocean biogeochemistry. Phil Trans Roy Soc B Biol Sci 375, 20190254 (2020).

26. L. J. Ustick, et al., Metagenomic analysis reveals global-scale patterns of ocean nutrient limitation. Science 372, 287–291 (2021).

27. A. C. Martiny, M. L. Coleman, S. W. Chisholm, Phosphate acquisition genes in Prochlorococcus ecotypes: Evidence for genome-wide adaptation. Proc Natl Acad Sci USA 103, 12552–12557 (2006).

28. A. C. Martiny, Y. Huang, W. Li, Occurrence of phosphate acquisition genes in Prochlorococcus cells from different ocean regions. Environmental Microbiology 11, 1340– 1347 (2009).

29. L. Garczarek, et al., Cyanorak v2.1: a scalable information system dedicated to the visualization and expert curation of marine and brackish picocyanobacteria genomes. Nucl Acids Res 49, D667–D676 (2021).

30. A. G. Kent, C. L. Dupont, S. Yooseph, A. C. Martiny, Global biogeography of Prochlorococcus genome diversity in the surface ocean. The ISME Journal 10, 1856–1865 (2016).

31. Q. Song, A. L. Gordon, M. Visbeck, Spreading of the Indonesian throughflow in the Indian Ocean. J Phys Oceanogr 34, 772–792 (2004).

32. N. J. West, P. Lebaron, P. G. Strutton, M. T. Suzuki, A novel clade of Prochlorococcus found in high nutrient low chlorophyll waters in the South and Equatorial Pacific Ocean. ISME J 5, 933–944 (2011).

33. D. B. Rusch, A. C. Martiny, C. L. Dupont, A. L. Halpern, J. C. Venter, Characterization of Prochlorococcus clades from iron-depleted oceanic regions. Proc Natl Acad Sci USA 107, 16184–16189 (2010).

34. A. C. Martiny, S. Kathuria, P. M. Berube, Widespread metabolic potential for nitrite and nitrate assimilation among Prochlorococcus ecotypes. Proc Natl Acad Sci USA 106, 10787– 10792 (2009).

35. P. M. Berube, A. Rasmussen, R. Braakman, R. Stepanauskas, S. W. Chisholm, Emergence of trait variability through the lens of nitrogen assimilation in Prochlorococcus. eLife 8, e41043–e41043 (2019).

36. P. M. Berube, et al., Physiology and evolution of nitrate acquisition in Prochlorococcus. ISME J (2015) https://doi.org/10.1038/ismej.2014.211.

37. M. Burnat, B. Li, S. H. Kim, A. J. Michael, E. Flores, Homospermidine biosynthesis in the cyanobacterium Anabaena requires a deoxyhypusine synthase homologue and is essential for normal diazotrophic growth. Mol Microbiol 109, 763–780 (2018).

38. B. Wang, et al., A guanidine-degrading enzyme controls genomic stability of ethylene-producing cyanobacteria. Nat Commun 12, 5150 (2021).

39. J. W. Nelson, R. M. Atilho, M. E. Sherlock, R. B. Stockbridge, R. R. Breaker, Metabolism of free guanidine in Bacteria is regulated by a widespread riboswitch class. Mol Cell 65, 220–230 (2017).

40. P. S. Novichkov, et al., RegPrecise 3.0 – A resource for genome-scale exploration of transcriptional regulation in bacteria. BMC Genomics 14, 745 (2013).

41. N. A. Kamennaya, A. F. Post, Characterization of cyanate metabolism in marine Synechococcus and Prochlorococcus spp. Appl Environ Microbiol 77, 291–301 (2011).

42. N. A. Kamennaya, A. F. Post, Distribution and expression of the cyanate acquisition potential among cyanobacterial populations in oligotrophic marine waters. Limnol Oceanogr 58, 1959–1971 (2013).

43. L. B. Jones, P. Ghosh, J.-H. Lee, C.-N. Chou, D. A. Y. 2018 Kunz, Linkage of the Nit1C gene cluster to bacterial cyanide assimilation as a nitrogen source. Microbiol 164, 956–968 (2018).

44. J. K. Saunders, G. Rocap, Genomic potential for arsenic efflux and methylation varies among global Prochlorococcus populations. ISME J 10, 197–209 (2016).

45. G. F. Riedel, Influence of salinity and sulfate on the toxicity of chromium(vi) to the estuarine diatom Thalassiosira Pseudonana. Journal of Phycology 20, 496–500 (1984).

46. F. Pablo, J. L. Stauber, R. T. Buckney, Toxicity of cyanide and cyanide complexes to the marine diatom Nitzschia closterium. Water Res 31, 2435–2442 (1997).

47. R. Pernil, S. Picossi, V. Mariscal, A. Herrero, E. Flores, ABC-type amino acid uptake transporters Bgt and N-II of Anabaena sp. strain PCC 7120 share an ATPase subunit and are expressed in vegetative cells and heterocysts. Mol Microbiol 67, 1067–1080 (2008).

48. M. L. Coleman, et al., Genomic islands and the ecology and evolution of Prochlorococcus. Science 311, 1768–1770 (2006).

49. C. M. Moore, et al., Processes and patterns of oceanic nutrient limitation. Nature Geosci 6, 701–710 (2013).

50. S. G. Tetu, et al., Microarray analysis of phosphate regulation in the marine cyanobacterium Synechococcus sp. WH8102. ISME J 3, 835–849 (2009).

51. L. L. Clark, E. D. Ingall, R. Benner, Marine phosphorus is selectively remineralized. Nature 393, 426–426 (1998).

52. R. Feingersch, et al., Potential for phosphite and phosphonate utilization by Prochlorococcus. ISME J 6, 827–834 (2012).

53. A. Martinez, G. W. Tyson, E. F. Delong, Widespread known and novel phosphonate utilization pathways in marine bacteria revealed by functional screening and metagenomic analyses. Environ Microbiol 12, 222–238 (2010).

54. O. A. Sosa, J. R. Casey, D. M. Karl, Methylphosphonate oxidation in Prochlorococcus strain MIT9301 supports phosphate acquisition, formate excretion, and carbon assimilation into purines. Appl Environ Microbiol 85, e00289–19 (2019).

55. A. Martínez, M. S. Osburne, A. K. Sharma, E. F. DeLong, S. W. Chisholm, Phosphite utilization by the marine picocyanobacterium Prochlorococcus MIT9301. Environ Microbiol 14, 1363–1377 (2012).

56. F. R. McSorley, et al., PhnY and PhnZ comprise a new oxidative pathway for enzymatic cleavage of a carbon–phosphorus bond. J Am Chem Soc 134, 8364–8367 (2012).

57. R. R. Malmstrom, et al., Ecology of uncultured Prochlorococcus clades revealed through single-cell genomics and biogeographic analysis. ISME J 7, 184–198 (2013).

58. J. W. Cooley, W. F. J. Vermaas, Succinate dehydrogenase and other respiratory pathways in thylakoid membranes of Synechocystis sp. strain PCC 6803: capacity comparisons and physiological function. J Bacteriol (2001) (January 27, 2022).

59. C. Whitfield, Biosynthesis and assembly of capsular polysaccharides in Escherichia coli. Annu Rev Biochem 75, 39–68 (2006).

60. A. Buffet, E. P. C. Rocha, O. Rendueles, Nutrient conditions are primary drivers of bacterial capsule maintenance in Klebsiella. Proc Roy Soc B Biol Sci 288, 20202876 (2021).

61. D. J. Scanlan, et al., Ecological genomics of marine picocyanobacteria. Microbiol Mol Biol Rev 73, 249–299 (2009).

62. F. Dempwolff, H. M. Wischhusen, M. Specht, P. L. Graumann, The deletion of bacterial dynamin and flotillin genes results in pleiotrophic effects on cell division, cell growth and in cell shape maintenance. BMC Microbiol 12, 298 (2012).

63. D. J. Lea-Smith, et al., Thylakoid terminal oxidases are essential for the cyanobacterium Synechocystis sp. PCC 6803 to survive rapidly changing light intensities. Plant Physiol 162, 484–495 (2013).

64. D. J. Lea-Smith, P. Bombelli, R. Vasudevan, C. J. Howe, Photosynthetic, respiratory and extracellular electron transport pathways in cyanobacteria. Biochim Biophys Acta Bioenerget 1857, 247–255 (2016).

65. C. Kranzler, et al., Coordinated transporter activity shapes high-affinity iron acquisition in cyanobacteria. ISME J 8, 409–417 (2014).

66. C. Boulay, A. Wilson, S. D’Haene, D. Kirilovsky, Identification of a protein required for recovery of full antenna capacity in OCP-related photoprotective mechanism in cyanobacteria. Proc Natl Acad Sci USA 107, 11620–11625 (2010).

67. C. Six, M. Ratin, D. Marie, E. Corre, Marine Synechococcus picocyanobacteria: Light utilization across latitudes. Proc Natl Acad Sci USA 118 (2021).

68. X. Xia, et al., Phylogeography and pigment type diversity of Synechococcus cyanobacteria in surface waters of the northwestern Pacific Ocean. Environ Microbiol 19, 142–158 (2017).

69. T. Grébert, et al., Light color acclimation is a key process in the global ocean distribution of Synechococcus cyanobacteria. Proc Natl Acad Sci USA 115, E2010–E2019 (2018).

70. J. E. Sanfilippo, et al., Self-regulating genomic island encoding tandem regulators confers chromatic acclimation to marine Synechococcus. Proc Natl Acad Sci USA 113, 6077–6082 (2016).

71. M. Costa, et al., Structure of Hierridin C, synthesis of hierridins B and C, and evidence for prevalent alkylresorcinol biosynthesis in picocyanobacteria. J. Nat. Prod. 82, 393–402 (2019).

72. A. A. Larkin, A. C. Martiny, Microdiversity shapes the traits, niche space, and biogeography of microbial taxa: The ecological function of microdiversity. Environ Microbiol Rep 9, 55– 70 (2017).

73. S. Sunagawa, et al., Structure and function of the global ocean microbiome. Science 348, 1261359–1261359 (2015).

74. B. Zhang, S. Horvath, A general framework for weighted gene co-expression network analysis. Stat Appl Genet Mol Biol 4, Article17 (2005).

75. B. Szmrecsanyi, Grammatical Variation in British English Dialects: A Study in Corpus-Based Dialectometry (Cambridge University Press, 2012) https://doi.org/10.1017/CBO9780511763380.

76. Oksanen, J., et al., Vegan: Community Ecology Package. R package Version 2.4-3 (2017).

77. P. Langfelder, S. Horvath, WGCNA: an R package for weighted correlation network analysis. BMC Bioinfo 9, 559 (2008).

