## Supplemental Text and Figures for "Differential global distribution of marine picocyanobacteria gene clusters reveals distinct niche-related adaptive strategies"

**This PDF file includes:**

Supplementary text
Figures S1 to S18
Legends for Datasets S1 to S9
SI References

**Other supplementary materials for this manuscript include the following:**

Datasets S1 to S9

### Supplementary Information Text

### Description of picocyanobacterial WGCNA modules and correlations with environmental parameters and ESTUs.

#### Prochlorococcus *modules*

In order to better interpret the global distribution of picocyanobacterial gene content, gene modules obtained by WGCNA were correlated to the available environmental parameters (Figs. 2A-B, S1A-B) and the relative abundance of *Prochlorococcus* or *Synechococcus* ESTUs at each station (Fig. 2C-D, S1A-B). The brown module, corresponding to genes preferentially found in Fe-limited HNLC areas and strongly associated with the presence of HLIIIA, HLIVA and LLIB, is described in the main text. The blue module was found to be associated to warm, low-chlorophyll oligotrophic regions with low N and P concentrations and high Fe availability (Fig. 2A), where ESTUs HLIIA dominate the *Prochlorococcus* community and HLIIB-D were also present at lower abundance (Fig. 2C, Fig. S1A). The turquoise module seems to correspond to genes present in cold, chlorophyll-rich waters, colonized by LLIA ESTUs, and to a lower extent to LLIC and LLID, but anti-correlated with HLII ESTUs. The turquoise module gathers station TARA-070, where LLIA is dominating (Fig. 1A), as well as stations dominated either by HLIA or the coldest stations dominated by HLIIA ESTUs (TARA-0146 and 149), the common point between all these stations being a strong relative abundance of LLIA (Figs. 1 and S1A). At last, the red module seems to be characteristic of cold, Fe-rich, N- and P-depleted waters, and strongly correlated to HLIA and anti-correlated to HLII-IV and LLIB ESTUs (Fig. 2A-C), corresponding to assemblages mainly found at the highest latitude stations of the *Tara* Oceans transect (TARA_066, 068, 093, 094, 133, 150, 151, 152) as well at all stations of the Mediterranean Sea (Fig. S1A).

#### Synechococcus *modules*

The same analysis performed on *Synechococcus* genes shows that the yellow module is correlated to phosphate and ammonium concentrations and strongly anti-correlated to Fe availability, and thus corresponds to genes found in HNLC areas. Accordingly, this module is correlated to ESTUs CRD1A, CRD1C, EnvAA and EnvBA, previously reported to dwell in Fe-depleted areas ((1, 2); Figs. 2B-D). Although the midnightblue module is only positively correlated to oxygen concentration, it is most strongly associated with ESTU IA and IVA-C, known to colonize cold, coastal, or mixed open ocean waters at high latitude (2) and anti-correlated with ESTUs IIA and IIIA/B (Fig. 2C-D). In terms of distribution, genes of this module are only found in two upwelling stations (TARA_093, 133), as well as in a cold station sampled in winter at the northernmost Atlantic station of the *Tara* Ocean transect, TARA_152 (Figs. 1B, S1B in this study and Fig.4 in (2)). The tan module was found in cold, chlorophyll-rich waters with a high relative abundance of ESTUs IA and IVA-C (Fig. 2B-D) and was also detected in the most strongly mixed waters of the *Tara* Oceans dataset, notably the upwelling stations (TARA_067, 093), at TARA_145, a cold station sampled in winter, North of the Gulf stream as well as in northern Atlantic stations of the *Tara* Ocean transect (TARA_150, 151, 152, Fig. S1B). The purple module is found in waters with high salinity, iron-rich, P-depleted waters, and associated with IIIA/B, WPC1A and all SC 5.3 ESTUs, known to co-occur in low-P areas of the world ocean (Fig. 2B-D). Consistently, it was specifically found in Mediterranean Sea and the only station of the Gulf of Mexico (TARA_142, Fig. S1B). At last, the salmon module was associated to warm, Fe-rich waters. This module was most strongly associated to ESTU IIA and also to a lower extent to the fairly rare ESTUs VIIA and 5.3B and its eigengene accordingly has higher values at stations dominated by ESTU IIA (Fig. S1).

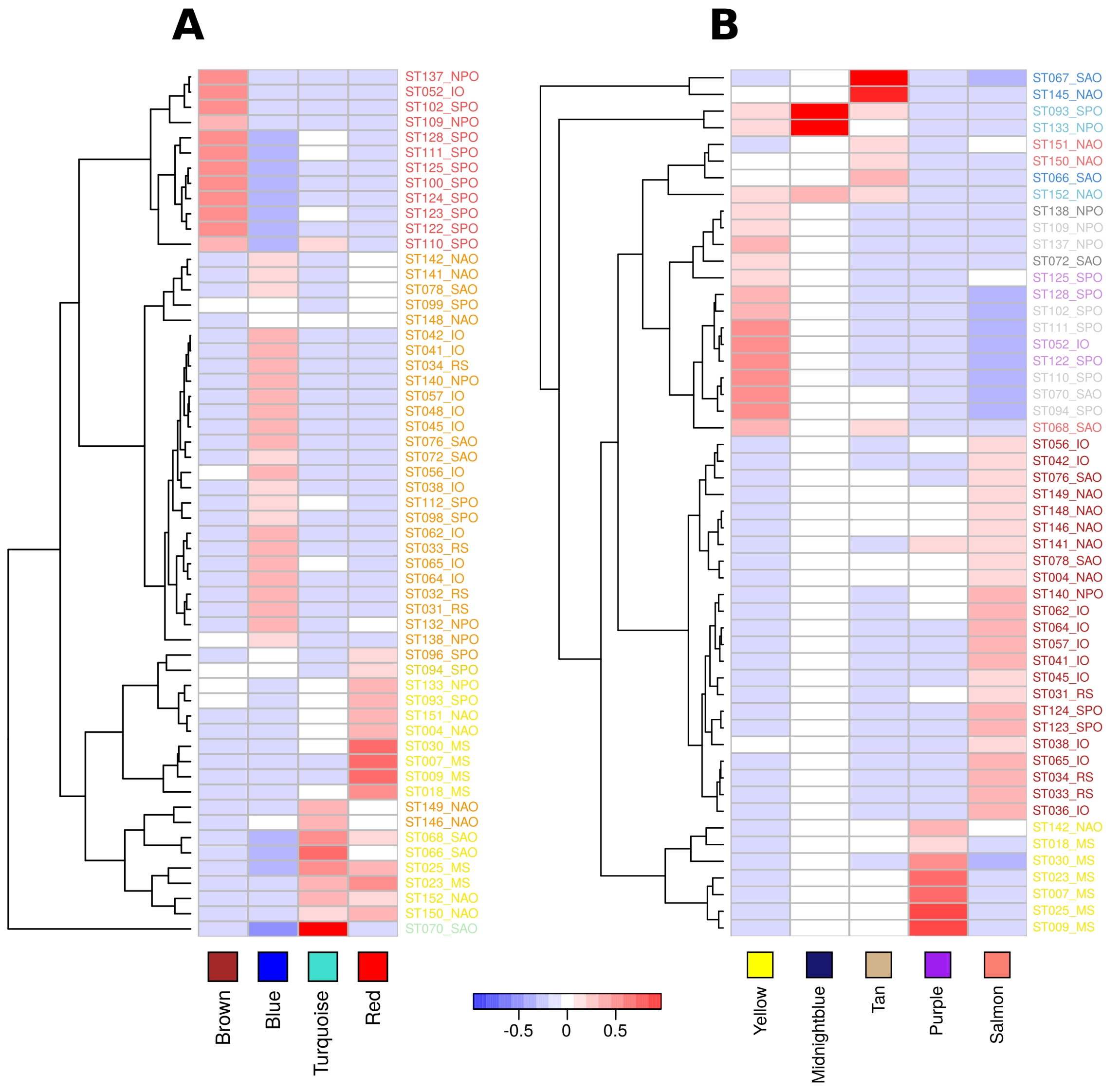


**Fig. S1. Distribution of the eigengene of each WGCNA module.** *Prochlorococcus* (*A*) and *Synechococcus* (*B*) modules are designated by color names indicated below each heatmap. The *eigengene* of a given module represents a consensus of the normalized relative abundance of genes of that module in *Tara* Oceans stations. Station names are colored according to ESTU assemblages defined in Farrant et al. (2016) and specify the oceanic region of each station as follows: SAO, South Atlantic Ocean; MS, Mediterranean Sea; NAO, North Atlantic Ocean; IO, Indian Ocean; RS, Red Sea; SPO, South Pacific Ocean; NPO, North Pacific Ocean.

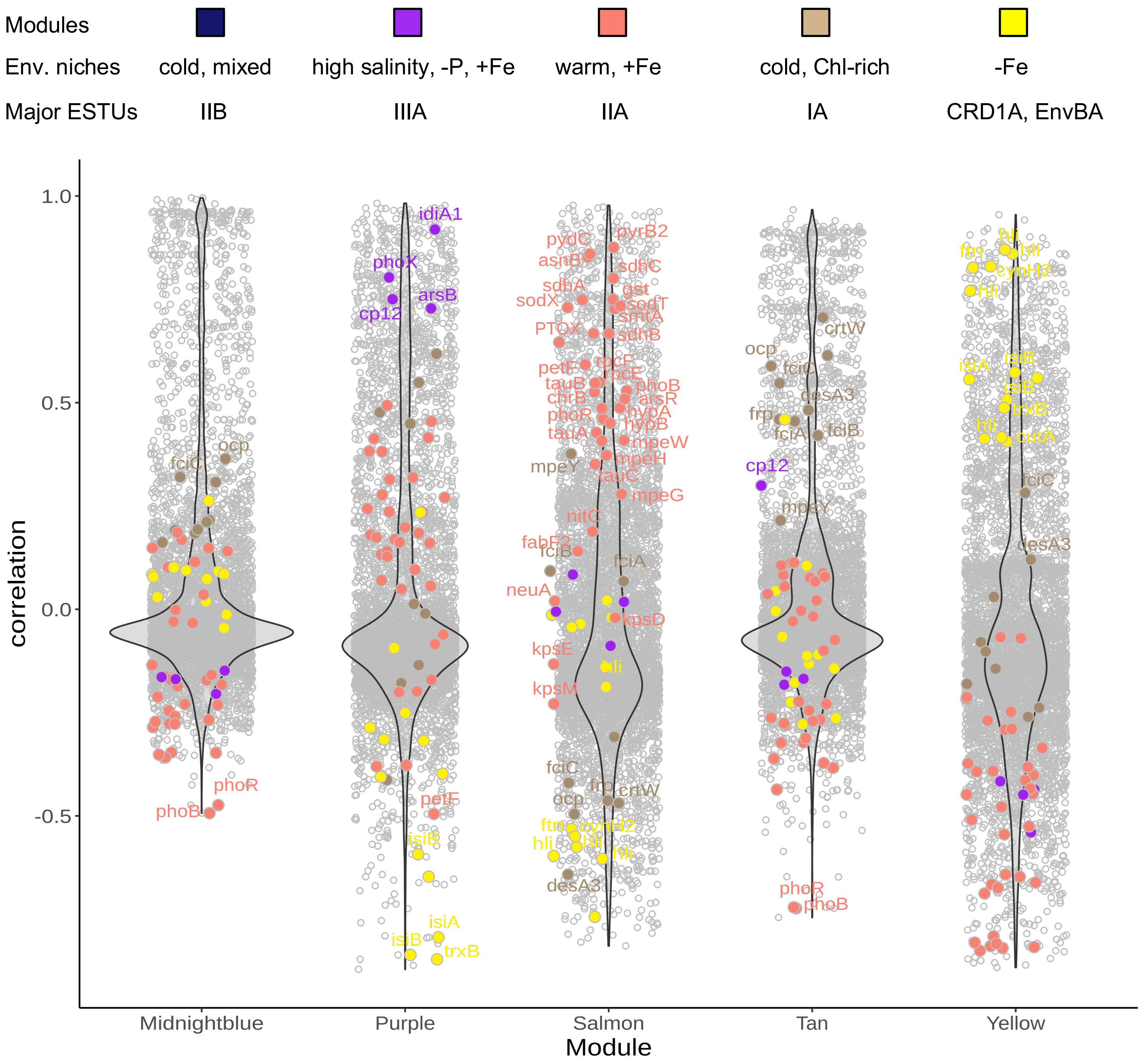


**Fig. S2. Same as Fig. 3 for *Synechococcus***

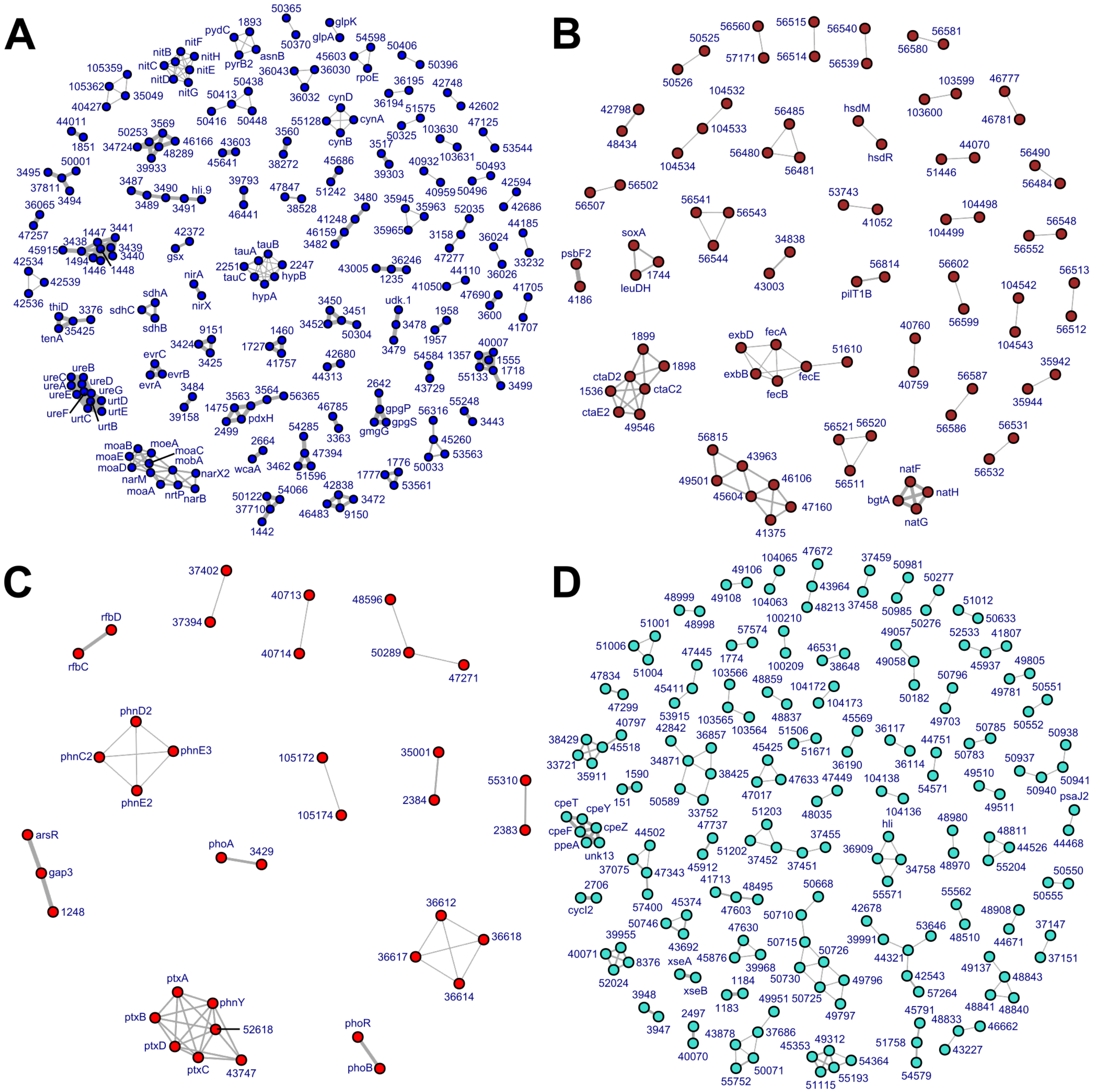


**Fig. S3. Same as Fig. 4 but for each individual *Prochlorococcus* WGCNA modules.** A link between two nodes indicates that these two genes are less than 5 genes apart in at least one genome and the thickness of this link is proportional to the number of genomes in which it is the case, with six distinct classes of link thickness: [1, 10[, [10, 20[, [20, 40[, [60, 80[ and [80, +100[ genomes..

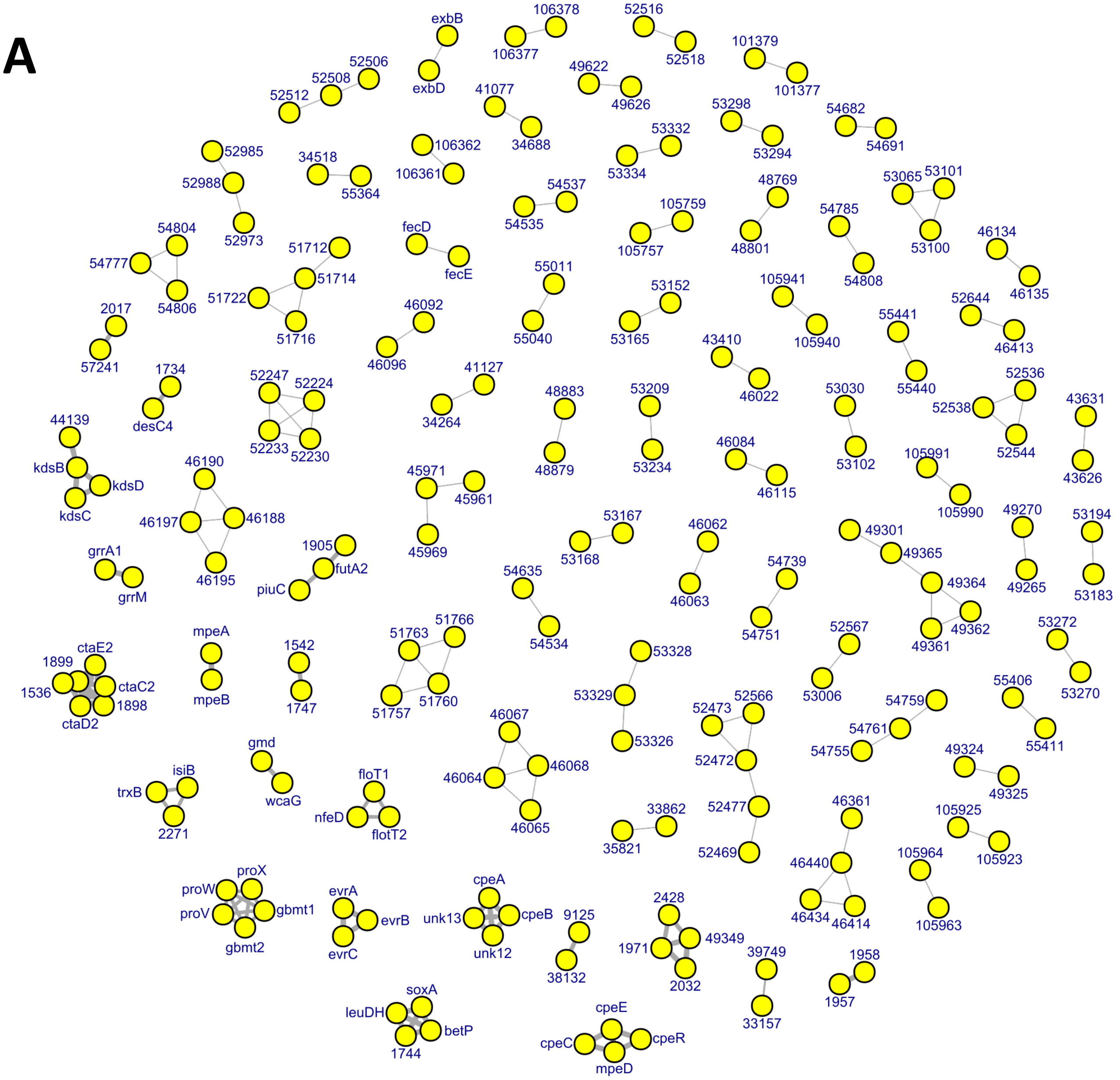


**Fig. S4. Same as Fig. S3 for *Synechococcus* yellow module (*A*)**

# B

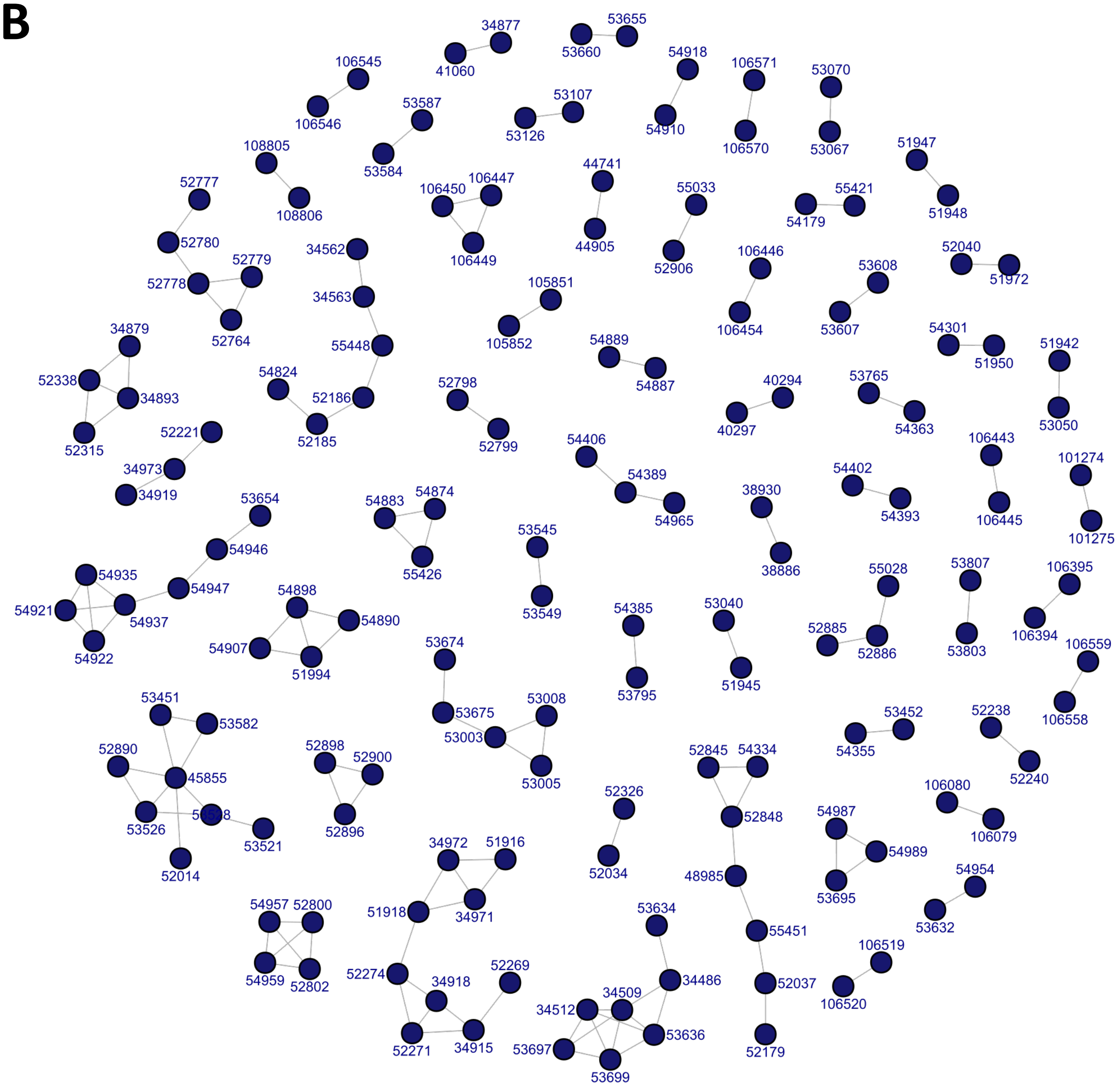


**Fig. S4. Continued for *Synechococcus* midnight blue module** (*B*)

**C**

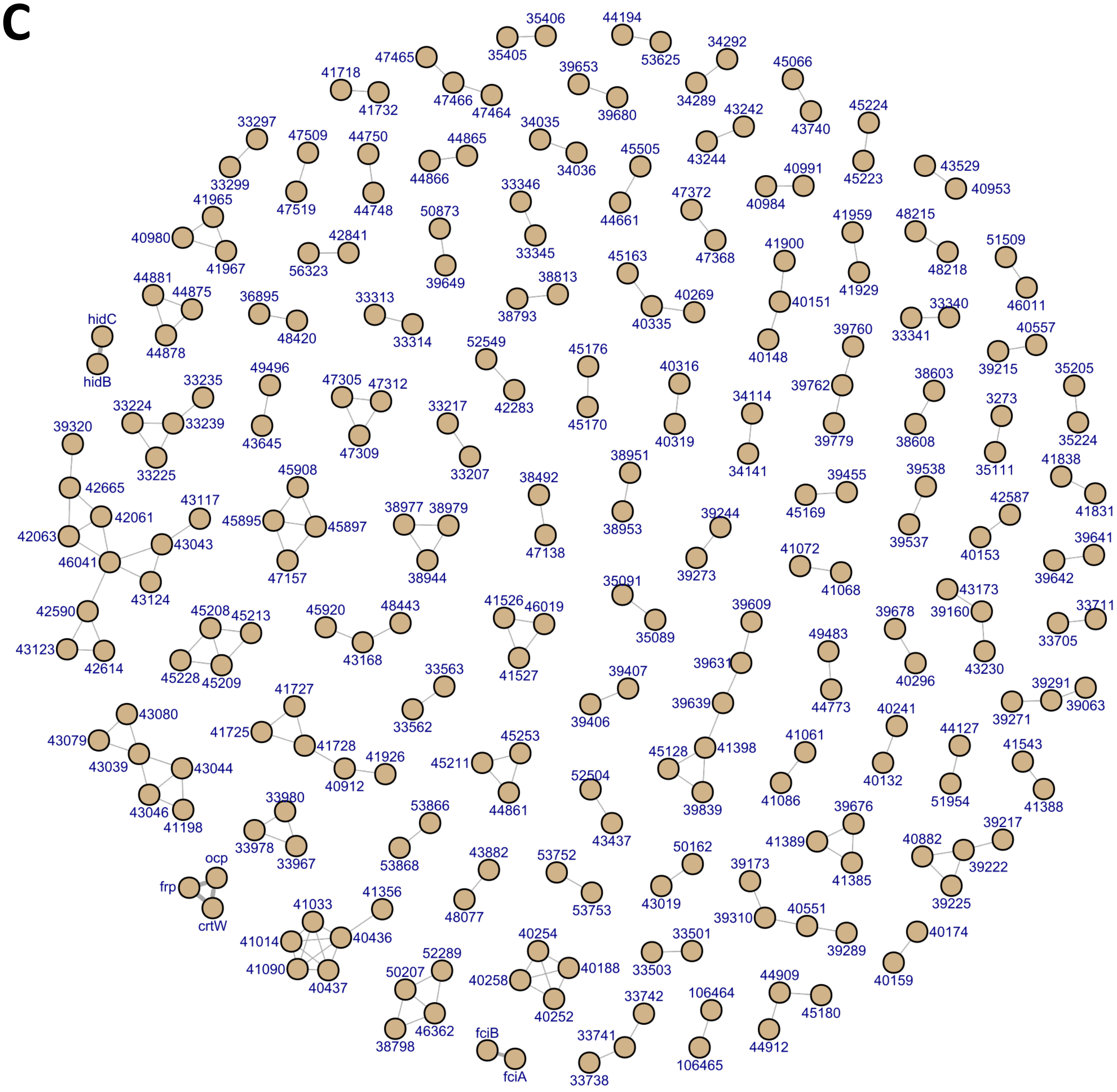


**Fig. S4. Continued for *Synechococcus* tan module (*C*)**

**D**

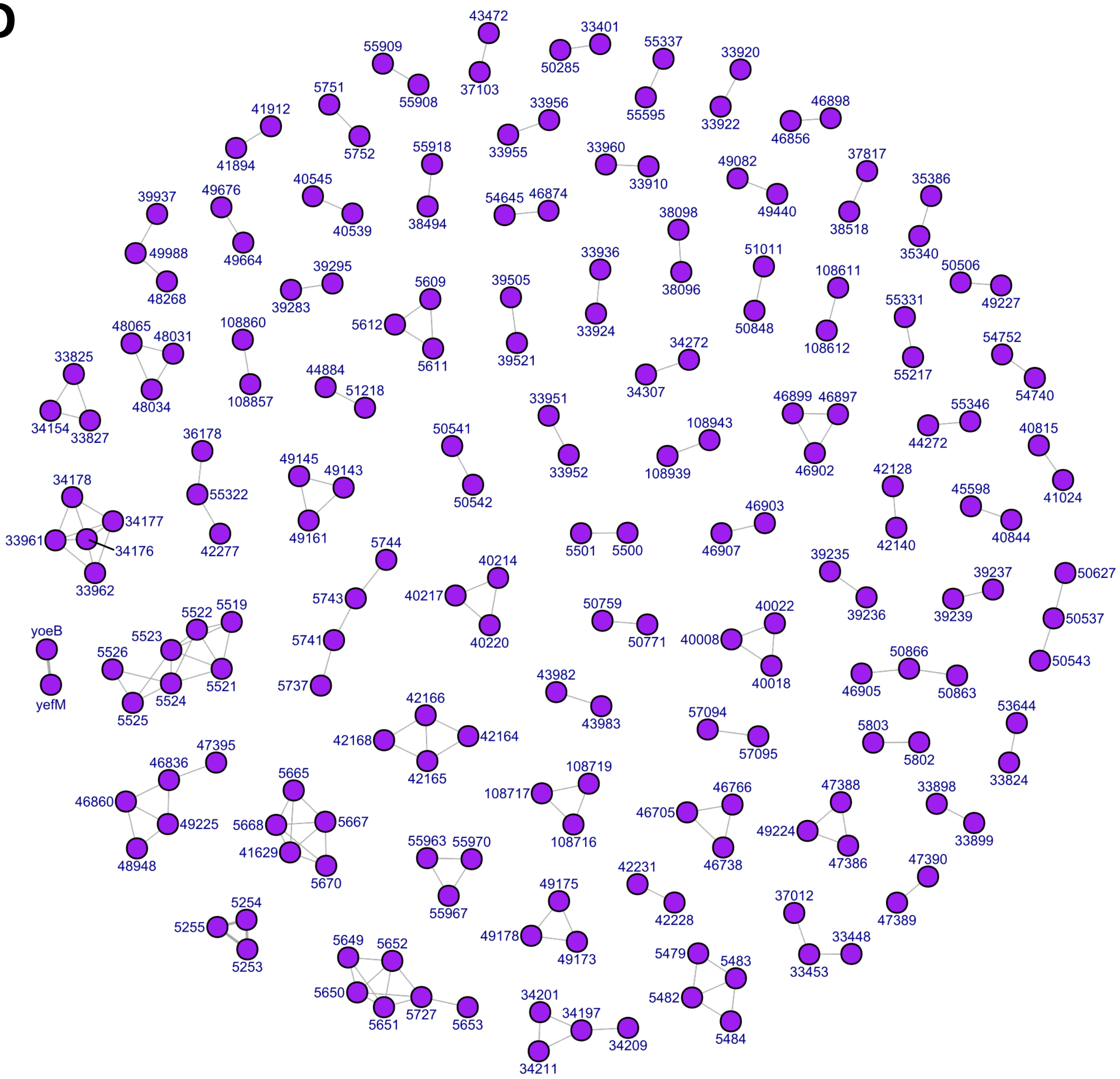


**Fig. S4. Continued for *Synechococcus* purple module** (*D*)

**E**

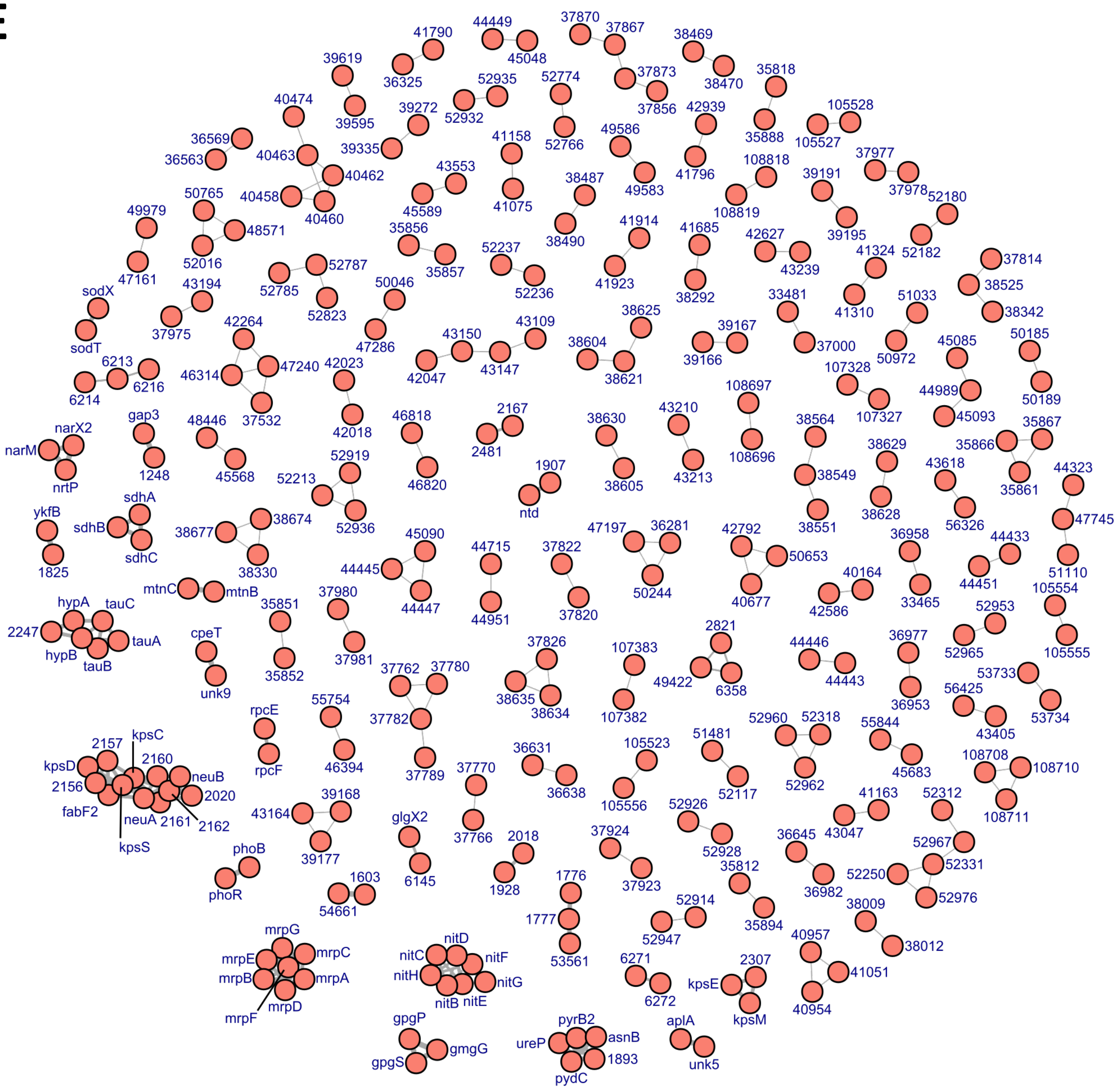


**Fig. S4. Continued for *Synechococcus* salmon module (*E*)**

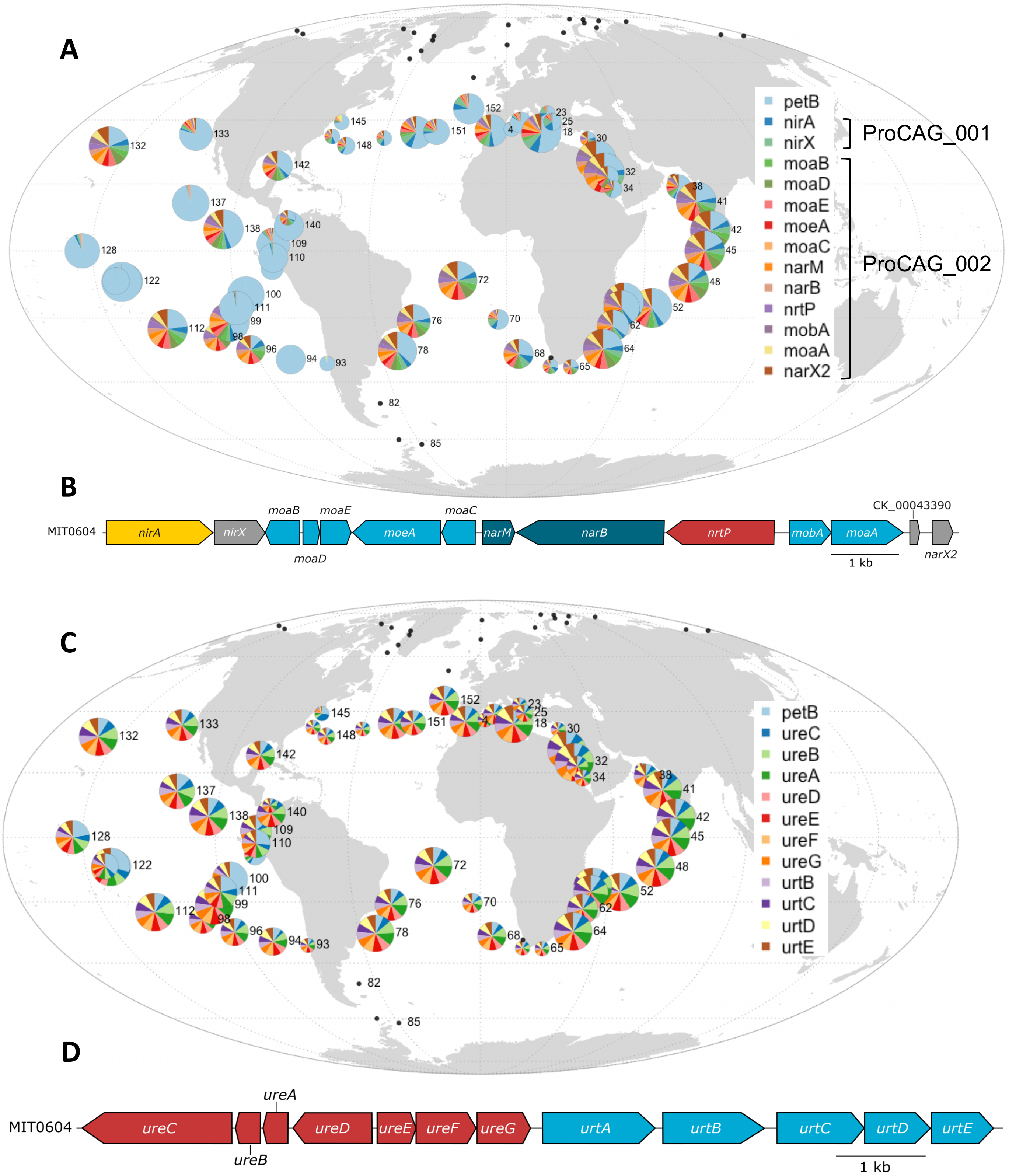


**Fig. S5. Global distribution map and genome organization of the *Prochlorococcus* CAGs involved in transport and assimilation of inorganic nitrogen and of urea.** The size of the circle is proportional to relative abundance of *Prochlorococcus* as estimated based on the single-copy core gene *petB* gene and this gene was also used to estimate the relative abundance of other genes in the population. (*A*) ProCAG_001 and 002 involved in transport and assimilation of inorganic nitrogen and (*B*) corresponding genomic region in *P. marinus* MIT0604. (*C*) ProCAG_003 involved in transport and assimilation of urea and (D) corresponding genomic region in *P. marinus* MIT0604

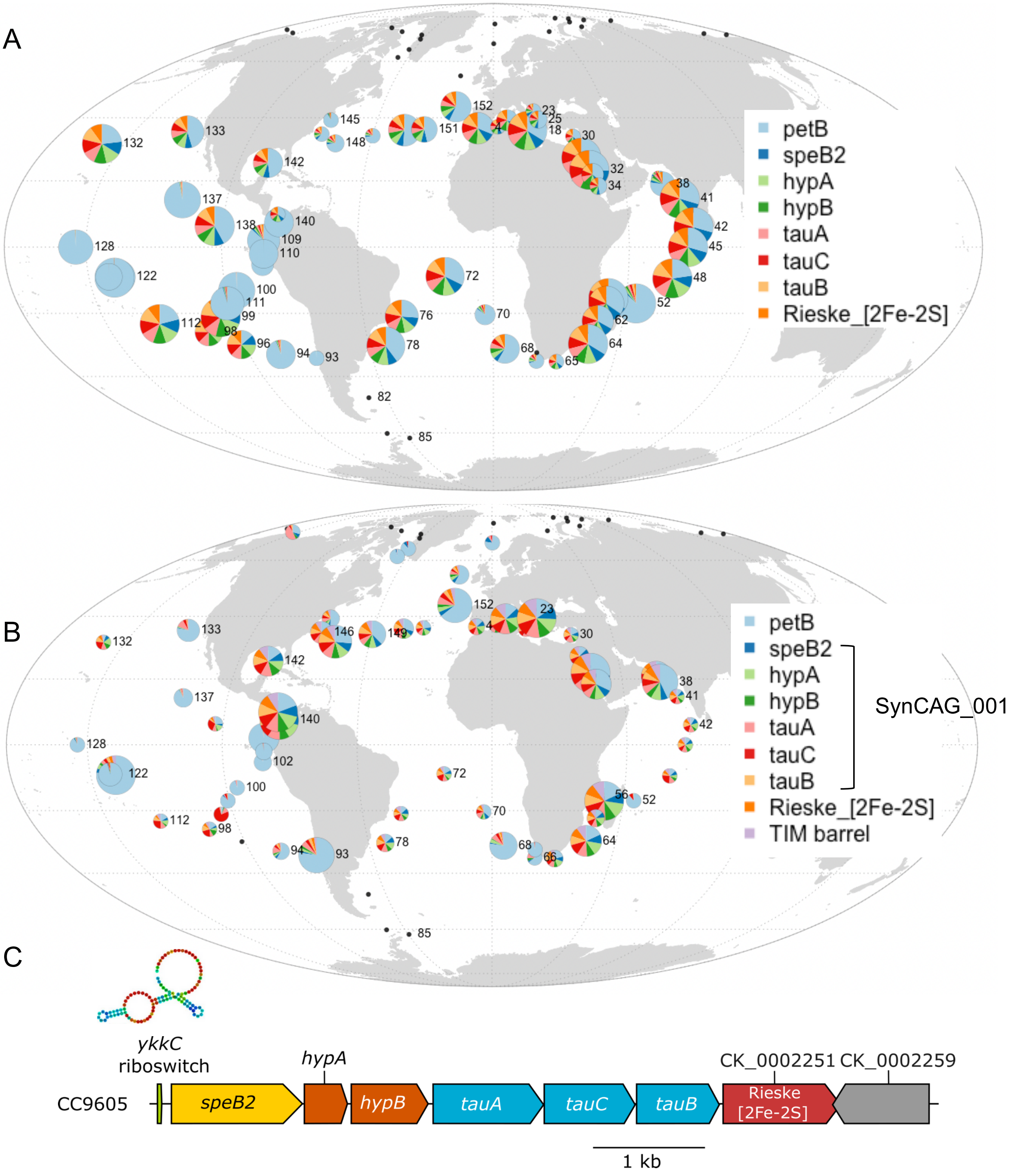


**Fig. S6. Global distribution map of the guanidinase CAG.** The size of the circle is proportional to relative abundance of each genus as estimated based on the single-copy core gene *petB* gene and this gene was also used to estimate the relative abundance of other genes in the population. **A.** *Prochlorococcus* ProCAG_004, **B.** *Synechococcus* SynCAG_001 as well as the CK_00002251 and CK_00002259, encoding an iron–sulfur protein and a TIM barrel domain-containing protein, respectively. Note that these two latter genes are not included in SynCAG_001 since they are absent from a few *Synechococcus/Cyanobium* genomes, see Dataset 6). **C.** Guanidinase gene cluster in *Synechococcus* sp. WH8102 starting with the *ykkC* riboswitch as predicted by regPrecise (https://regprecise.lbl.gov/regulon.jsp?regulon_id=23874)

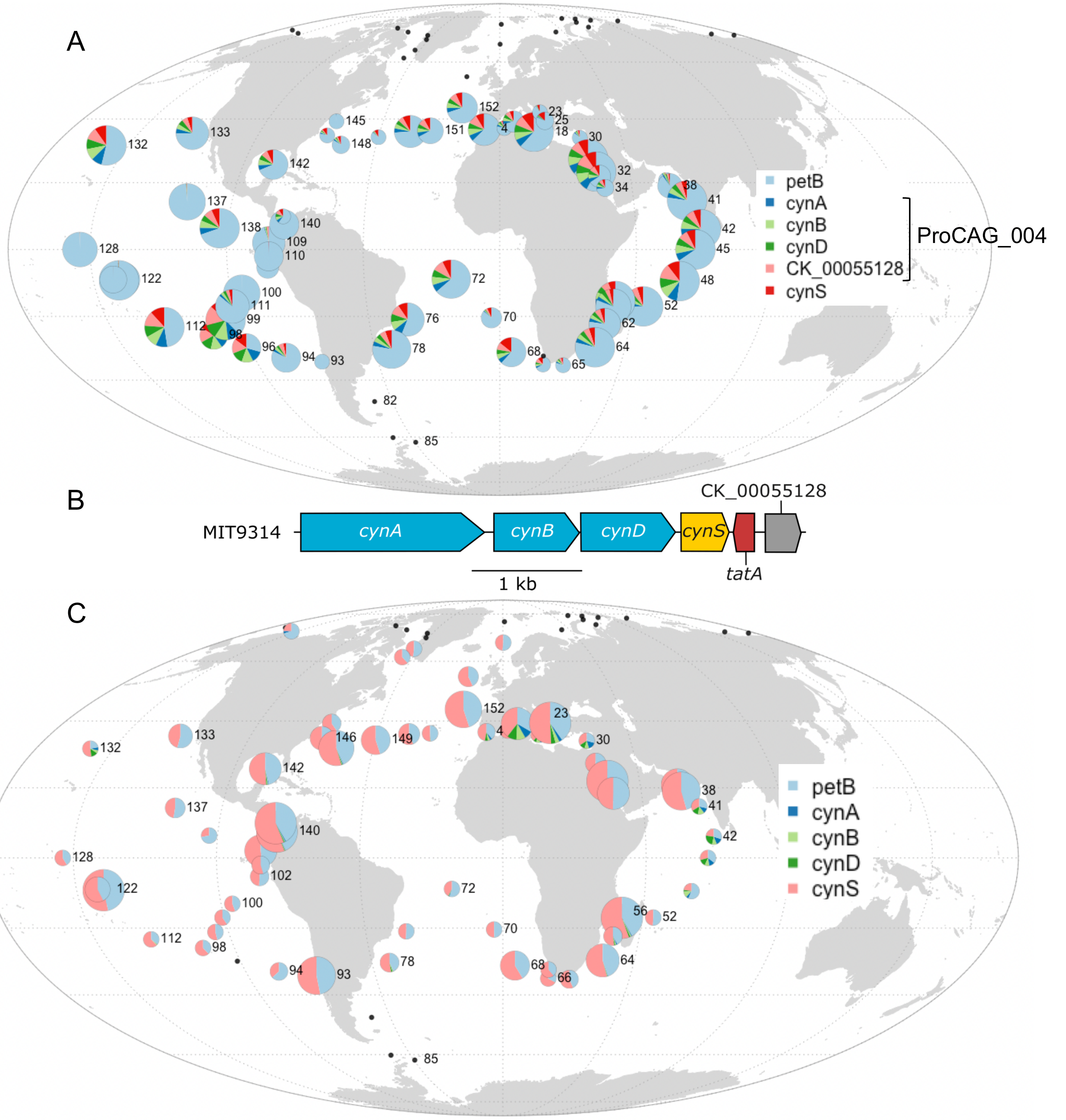


**Fig. S7. Global distribution map of picocyanobacterial CAGs involved in the uptake and degradation of cyanate.** The size of the circle is proportional to relative abundance of each genus as estimated based on the single-copy core gene *petB* gene and this gene was also used to estimate the relative abundance of other genes in the population. (*A*) *Prochlorococcus* CAG (ProCAG_005) involved in cyanate transport and uptake. Note that this CAG does not include *cynS* due its presence without *cynABD* in several LLI genomes. (*B*) Genomic region in *Prochlorococcus marinus* MIT9314. (*C*) distribution of the same non-CAG gene operon in *Synechococcus*. Note that CK_00055128 is absent in *Synechococcus/Cyanobium.*

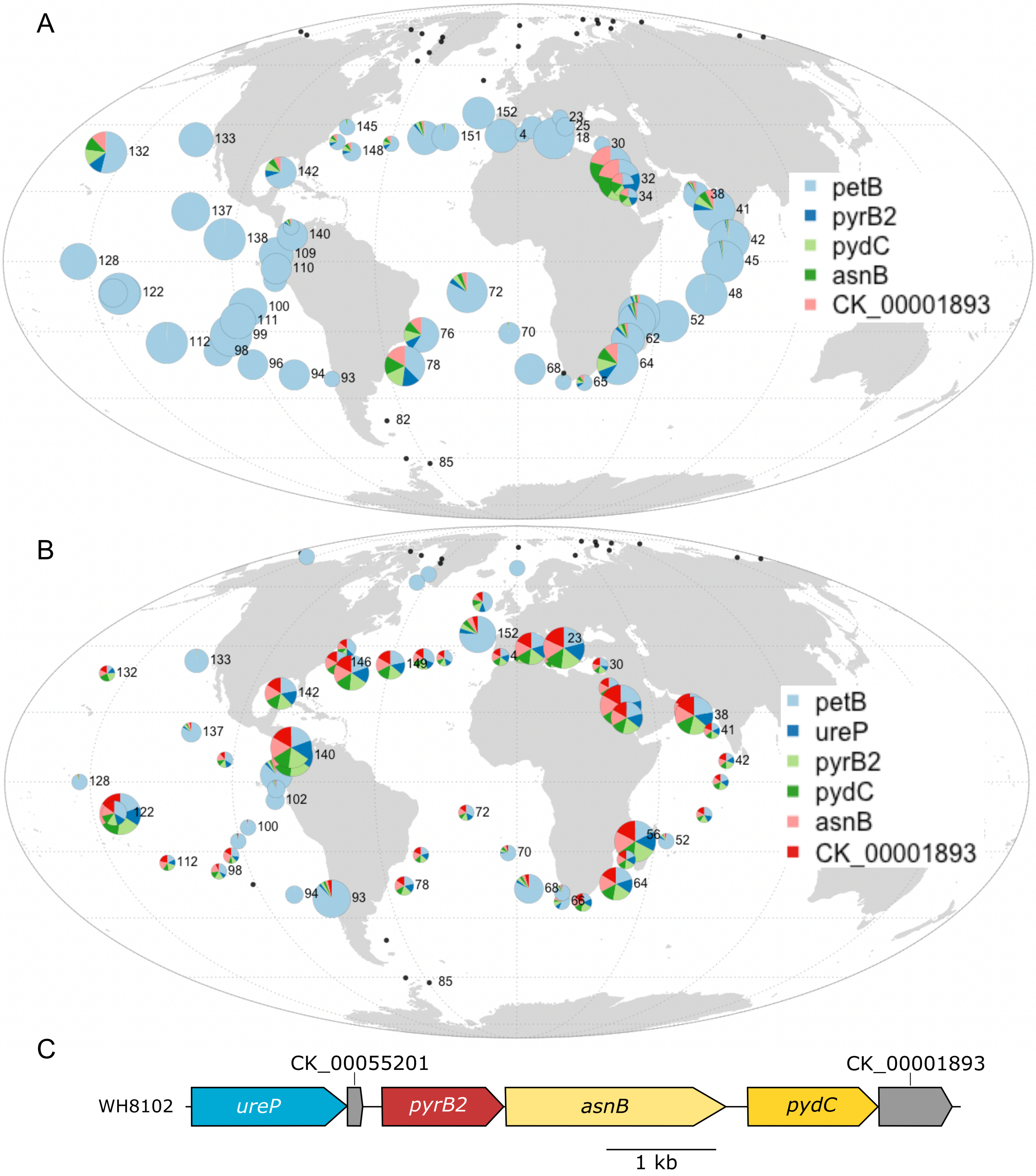


**Fig. S8. Global distribution map of picocyanobacterial CAGs involved in the biosynthesis of pyrimidines.** The size of the circle is proportional to relative abundance of each genus as estimated based on the single-copy core gene *petB* gene and this gene was also used to estimate the relative abundance of other genes in the population. CAG involved in the biosynthesis of pyrimidines in (*A*) *Prochlorococcus* (ProCAG_007) and (*B*) *Synechococcus* (SynCAG_003). (*C*) Genomic region in *Synechococcus* sp. WH8102.

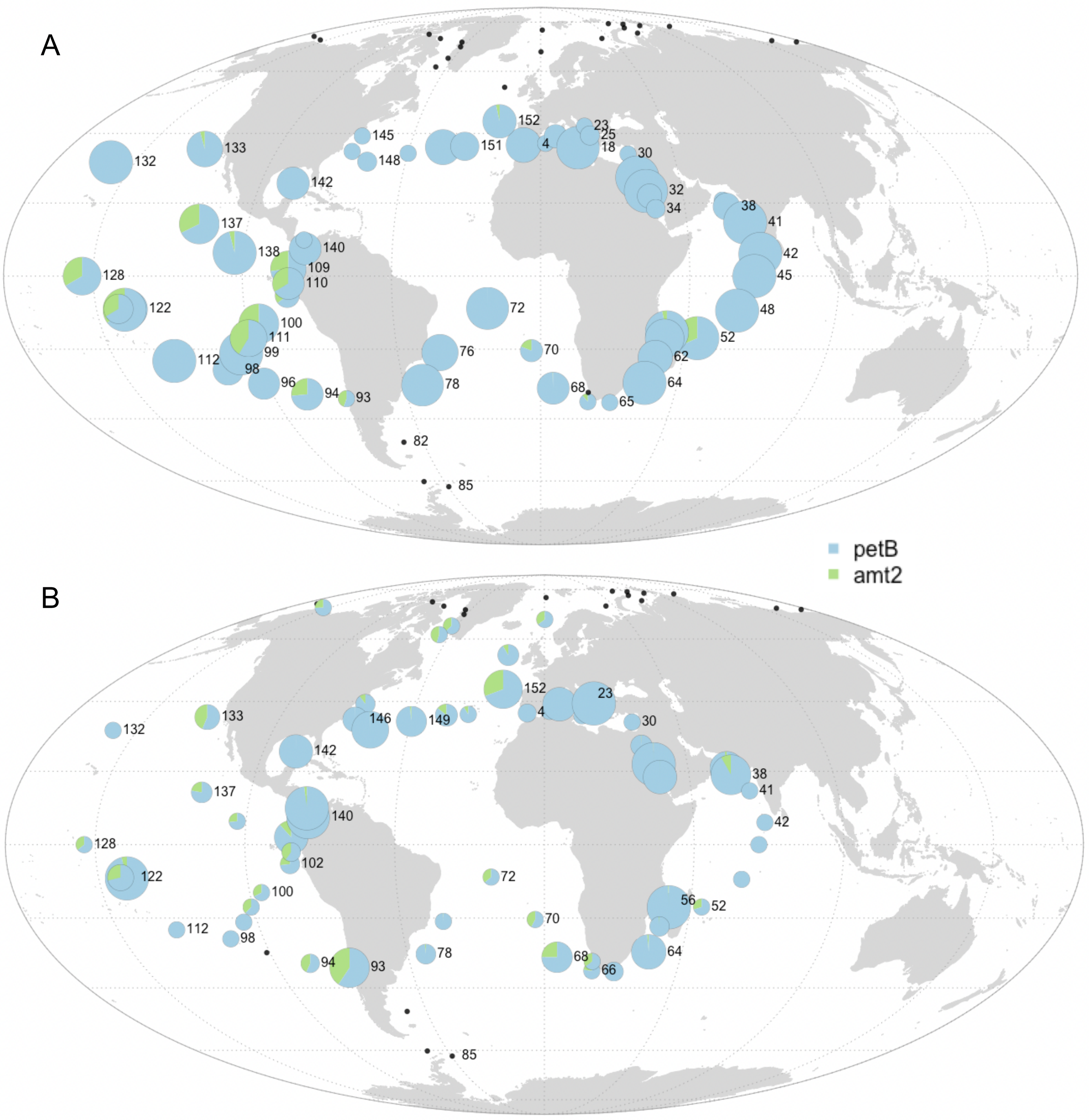


**Fig. S9. Global distribution map of the *amt2* gene, potentially involved in ammonium transport.** The size of the circle is proportional to relative abundance of each genus as estimated based on the single-copy core gene *petB* gene and this gene was also used to estimate the relative abundance of other genes in the population. (*A*) *Prochlorococcus*, (*B*) *Synechococcus.*

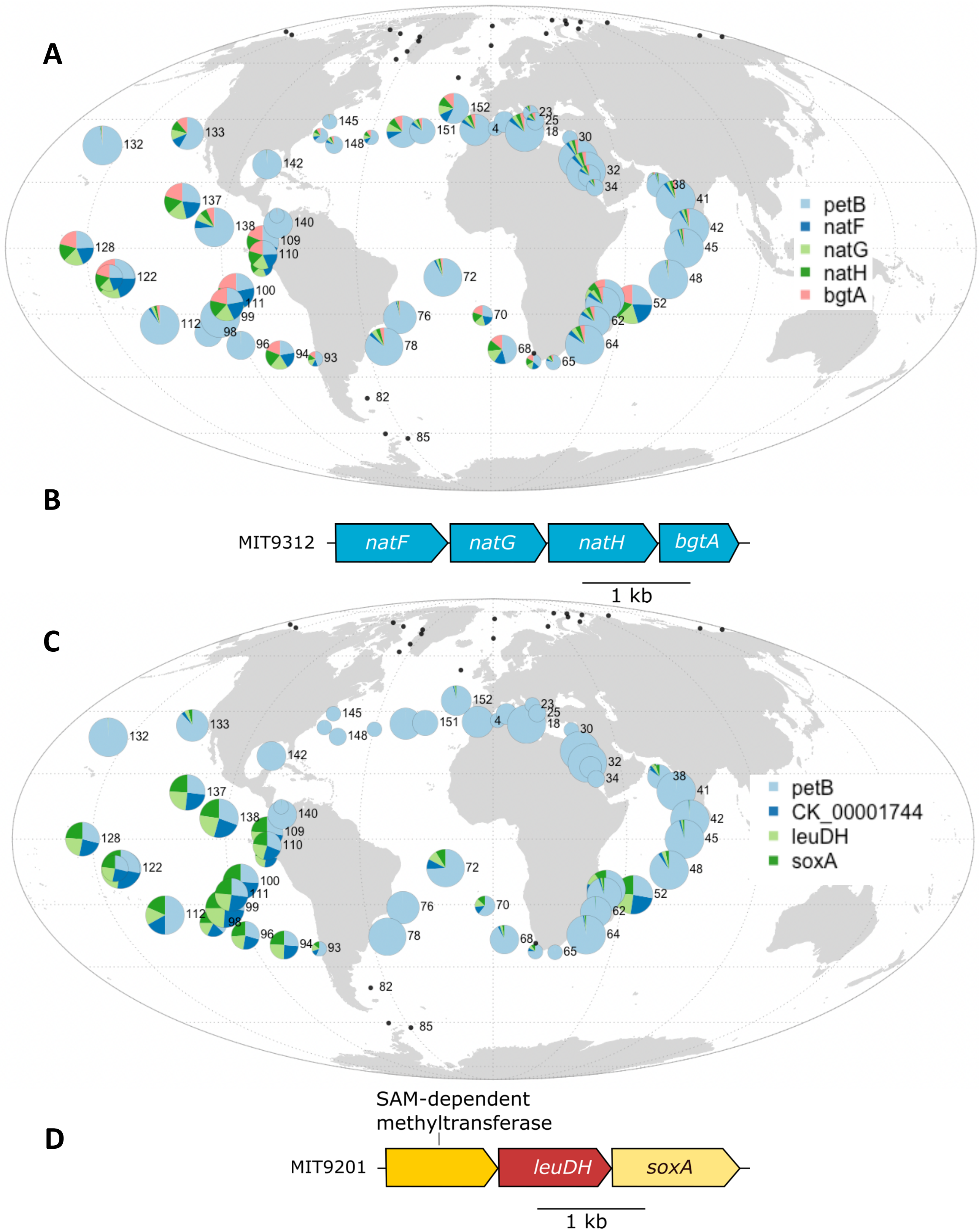


**Fig. S10. Global distribution map of the *Prochlorococcus* CAG putatively involved in amino acid transport or metabolism.** The size of the circle is proportional to relative abundance of *Prochlorococcus* as estimated based on the single-copy core gene *petB* gene and this gene was also used to estimate the relative abundance of other genes in the population. (*A*) CAG involved in ABC-type uptake transporter for acidic and neutral polar amino acids (ProCAG_008) and (*B*) corresponding genomic region in *P. marinus* MIT9312, (*C*) CAG putatively involved in amino acid metabolism (ProCAG_009) and (*D*) corresponding genomic region in *P. marinus* MIT9201

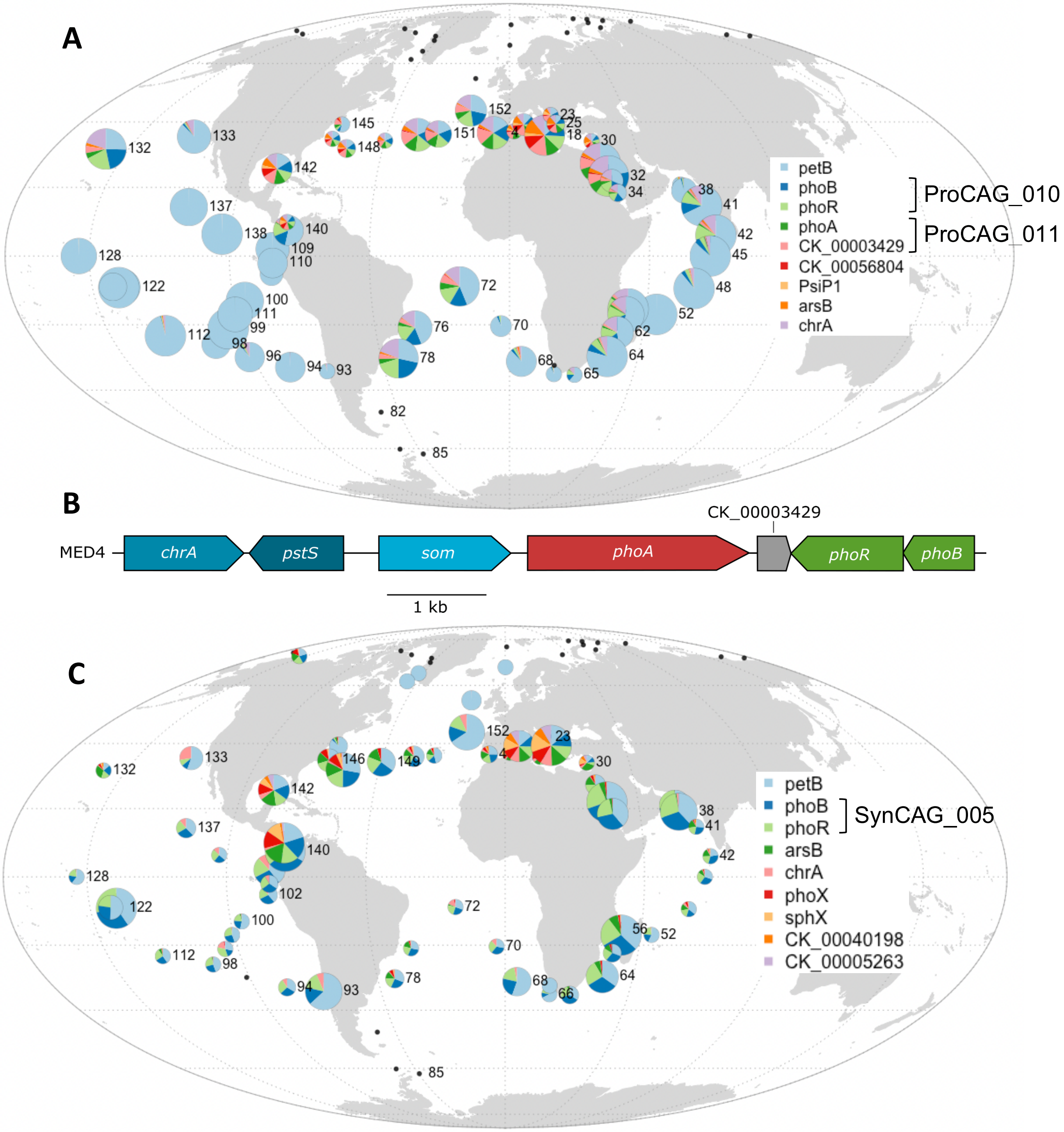


**Fig. S11. Global distribution map of picocyanobacterial genes putatively involved in adaptation to P-depletion.** The size of the circle is proportional to relative abundance of each genus as estimated based on the single-copy core gene *petB* gene and this gene was also used to estimate the relative abundance of other genes in the population. (*A*) *Prochlorococcus* ProCAG_010 and ProCAG_011 as well as genes often retrieved in the same genomic area. (*B*) and corresponding genomic region in *P. marinus* MED4. (*C*) *Synechococcus* SynCAG_005 and marker genes of P-limitation retrieved in the purple module, including CK_00040198 and CK_00052500 encoding putative alkaline phosphatases, both absent from reference *Prochlorococcus* genomes.

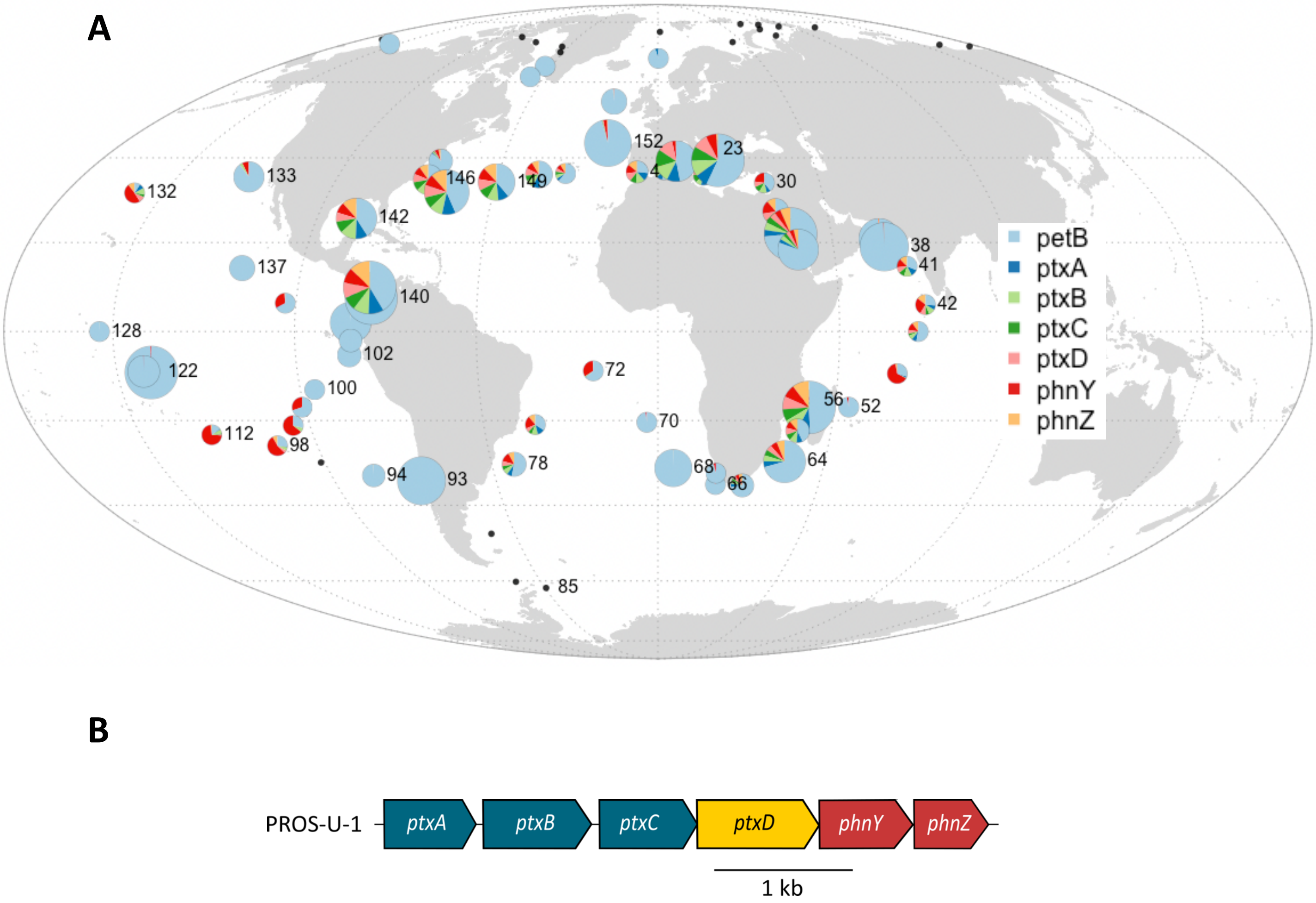


**Fig. S12. Global distribution map of *Synechococcus* genes potentially involved in phosphonate and phosphite transport and assimilation.** The size of the circle is proportional to relative abundance of each genus as estimated based on the single-copy core gene *petB* gene and this gene was also used to estimate the relative abundance of other genes in the population. (*A*) Global distribution map and (*B*) corresponding genomic region in *Synechococcus* sp. PROS-U-1.

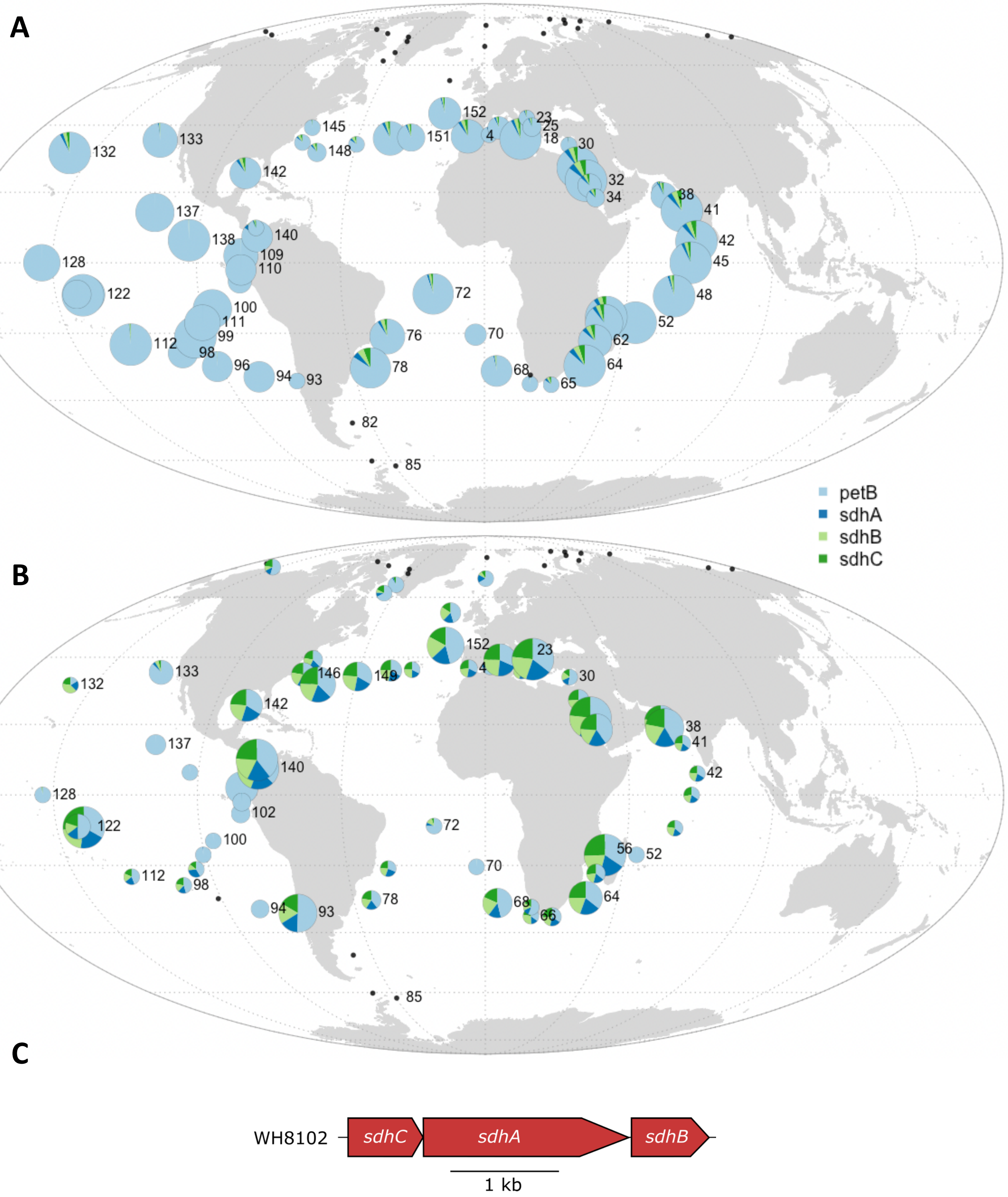


**Fig. S13. Global distribution map of CAGs involved in the biosynthesis of succinate deshydrogenase.** The size of the circle is proportional to relative abundance of each genus as estimated based on the single-copy core gene *petB* gene and this gene was also used to estimate the relative abundance of other genes in the population. (*A*) *Prochlorococcus* ProCAG_014, (*B*) *Synechococcus* SynCAG_006 and (*C*) corresponding genomic region in *Synechococcus* sp. WH8102.

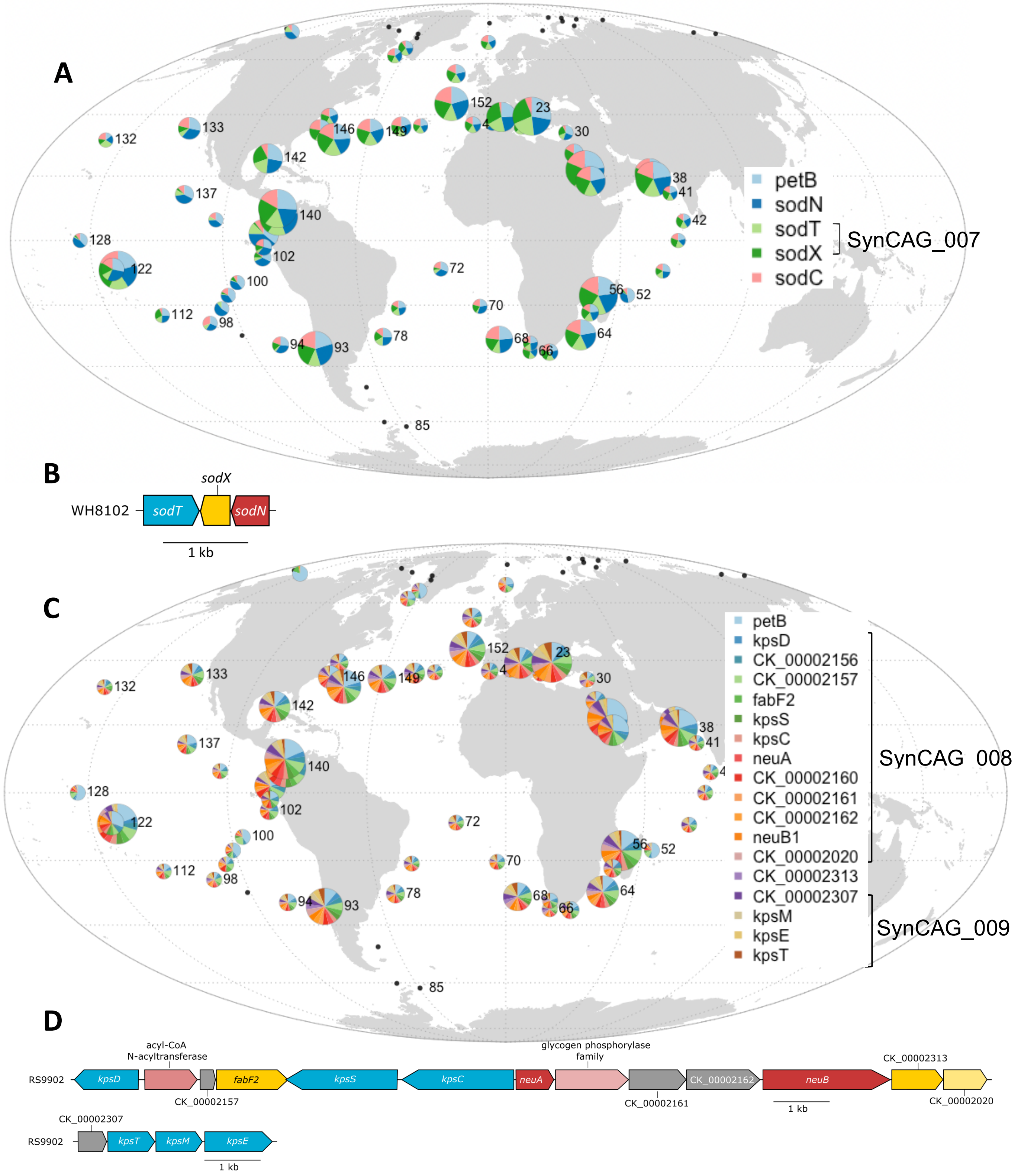


**Fig. S14. Global distribution map of *Synechococcus* CAGs specifically enriched in Fe-replete areas.** The size of the circle is proportional to relative abundance of each genus as estimated based on the single-copy core gene *petB* gene and this gene was also used to estimate the relative abundance of other genes in the population. (*A*) SynCAG_008 encompassing two genes related to nickel transport (*sodT*) and maturation (*sodX*) of the Ni-superoxide dismutase (*sodN*), the latter being also shown for comparison. (*B*) corresponding genomic region in *Synechococcus sp.* WH8102 (*C*) SynCAG_009 and other related gene putatively involved in the biosynthesis of polysaccharide capsules. (D) corresponding genomic region in *Synechococcus* sp. RS9902.

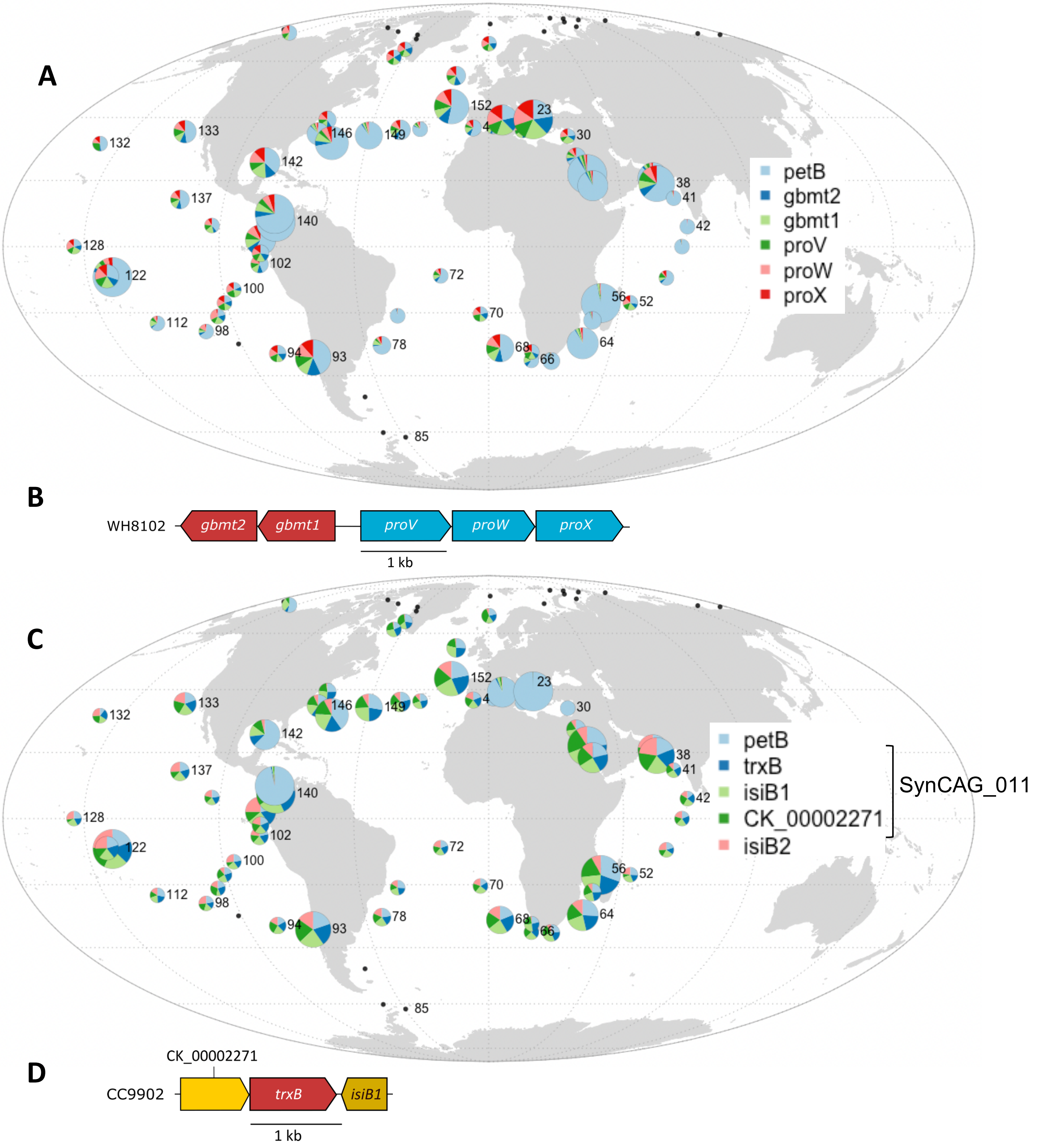


**Fig. S15. Global distribution map of the *Synechococcus* CAGs enriched in Fe-depleted areas.** The size of the circle is proportional to relative abundance of each genus as estimated based on the single-copy core gene *petB* gene and this gene was also used to estimate the relative abundance of other genes in the population. (A) SynCAG_010 involved in glycine betaine synthesis and transport and (*B*) its corresponding genomic region in *Synechococcus* sp. WH8102. (*C*) SynCAG_011 encoding a flavodoxin and a thioredoxin reductase and (*D*) its corresponding genomic region in *Synechococcus* sp. CC9902. The second *isiB* copy (*isiB2*) is shown here for comparison.

A

B

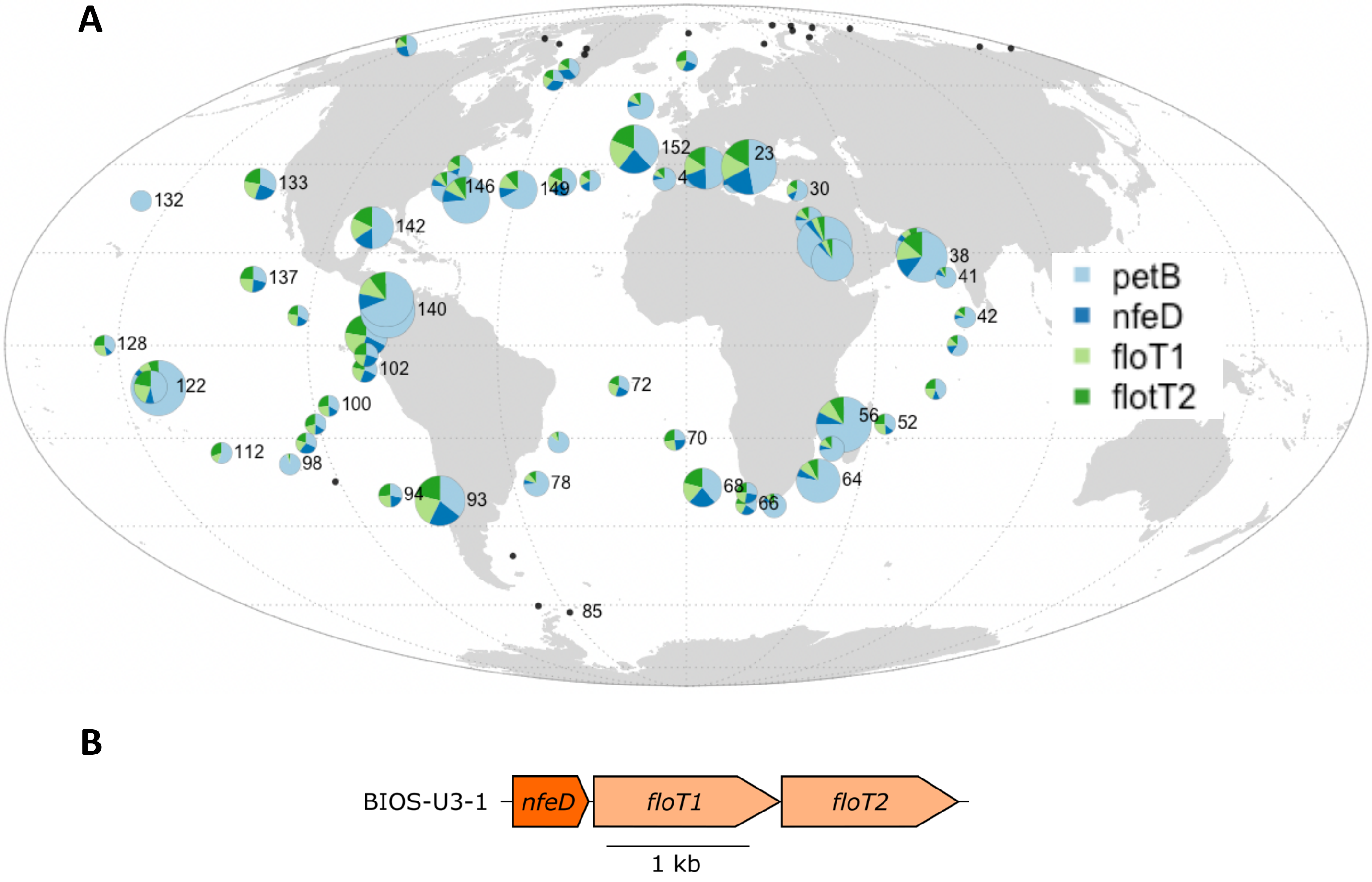


**Fig. S16. Global distribution map of the *Synechococcus* CAGs enriched in Fe-depleted areas (continued).** (*A*) SynCAG_012 involved in the production of lipid rafts and (*B*) its corresponding genomic region in *Synechococcus* sp. BIOS-U3-1**.** The size of the circle is proportional to relative abundance of each genus as estimated based on the single-copy core gene *petB* gene and this gene was also used to estimate the relative abundance of other genes in the population.

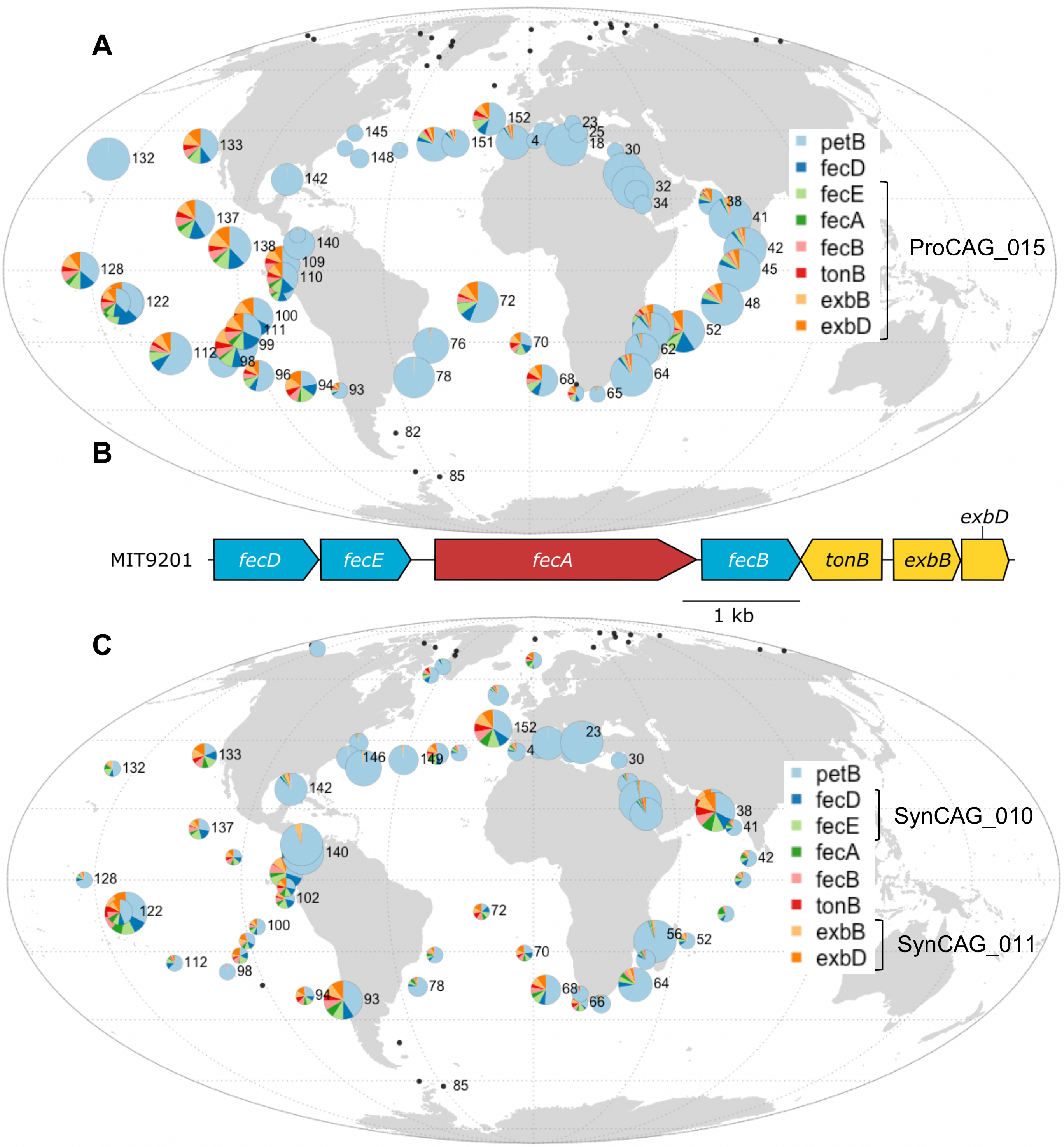


**Fig. S17. Global distribution map of CAGs involved in TonB-dependent siderophore uptake.** The size of the circle is proportional to relative abundance of each genus as estimated based on the single-copy core gene *petB* gene and this gene was also used to estimate the relative abundance of other genes in the population. (*A*) *Prochlorococcus* ProCAG_015 and (*B*) corresponding genomic region in *P. marinus* MIT9201. (*C*) *Synechococcus* SynCAG_010 and 011**.**

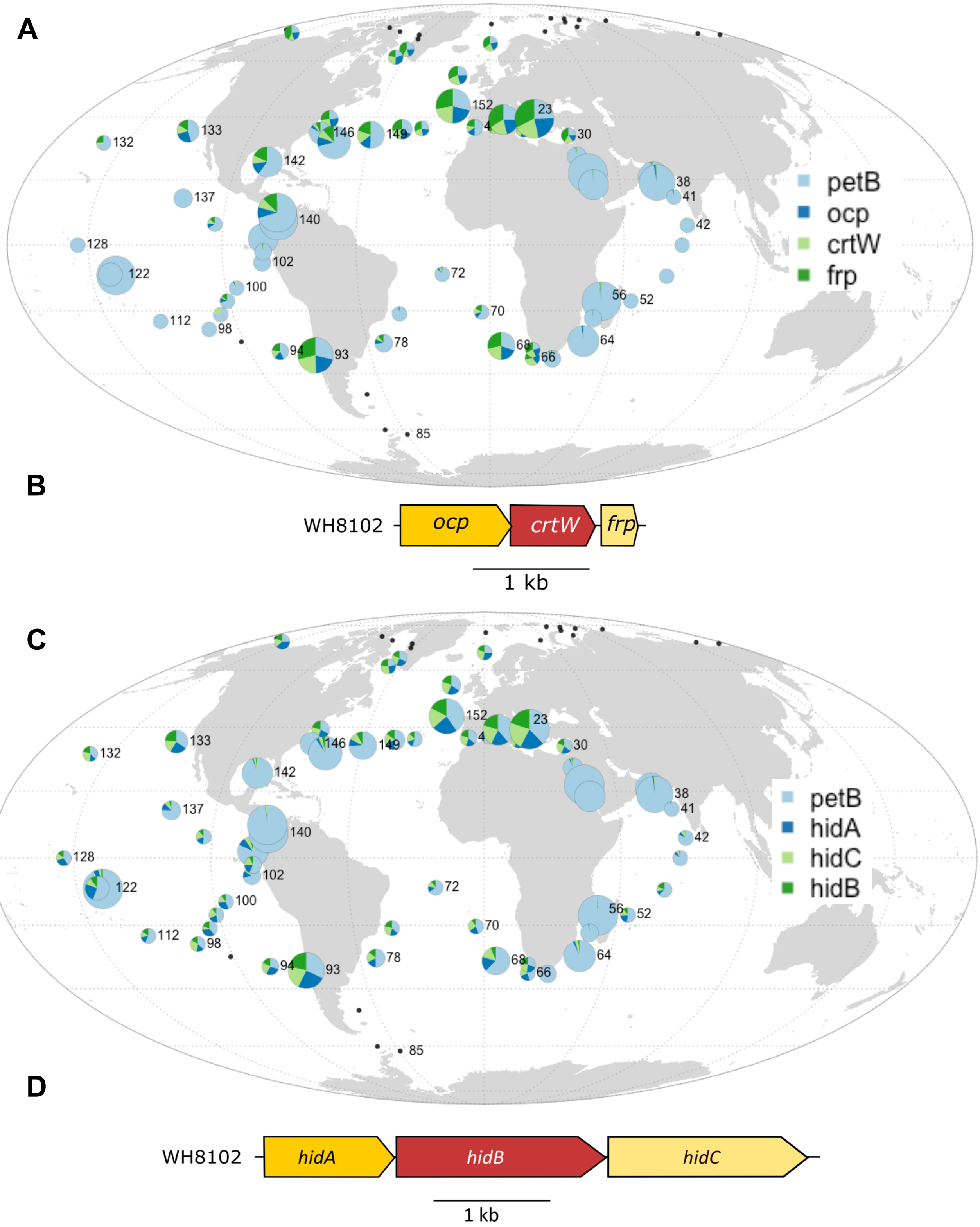


**Fig. S18. Global distribution map of the *Synechococcus* CAGs enriched in cold waters.** (*A*) SynCAG_016 involved in the orange caroteno-protein mediated photoprotection and (*B*) its corresponding genomic region in *Synechococcus* sp. WH8102. (C) SynCAG_017 involved in the biosynthesis of a hierridin C and (*D*) its corresponding genomic region in *Synechococcus* sp. WH8102. The size of the circle is proportional to relative abundance of *Synechococcus* as estimated based on the single-copy core gene *petB* gene and this gene was also used to estimate the relative abundance of other genes in the *Synechococcus* population.

**Dataset (separate file)**

**Dataset 1: Distance matrix based on relative abundances of *Synechococcus* CLOGs (Bray-Curtis distance)**

**Dataset 2: distance matrix based on relative abundances of *Synechococcus* ESTUs (Bray-Curtis distance)**

**Dataset 3: distance matrix based on relative abundances of *Prochlorococcus* CLOGs (Bray-Curtis distance)**

**Dataset 4: distance matrix based on relative abundances of *Prochlorococcus* ESTUs (Bray-Curtis distance)**

**Dataset 5: Inventory and description of all *Prochlorococcus* and *Synechococcus* CLOGs retrieved in WGCNA modules generated from picocyanobacterial reads extracted from *Tara* Oceans metagenomes.** Functional annotation and phyletic patterns of each CLOGs were derived from the Cyanorak v2.1 information system. Abbreviations: CLOG, clusters of likely orthologous genes; CK, Cyanorak; WGCNA, Weighted Correlation Network Analysis.

**Dataset 6: Inventory and description of *Prochlorococcus* and *Synechococcus* CLOGs retrieved in WGCNA modules and involved in CAGs or individual genes of interest mentioned in the text.** Functional annotation and phyletic patterns of each CLOG were derived from the Cyanorak v2.1 information system. Abbreviations: CLOG, clusters of likely orthologous genes; CK, Cyanorak; WGCNA, Weighted Correlation Network Analysis.

**Dataset 7: List of Cyanorak CLOGs belonging to clusters of adjacent genes for each *Prochlorococcus* WGCNA module**

**Dataset 8: List of Cyanorak CLOGs belonging to clusters of adjacent genes for each *Synechococcus* WGCNA module**

**Dataset 9: Genomes used in this study for whole genome recruitment, including outgroups.** The references are listed below the table.

**SI References**


1. Sohm, Jill A, Nathan A Ahlgren, Zachary J Thomson, Cheryl Williams, James W Moffett, Mak A Saito, Eric A Webb, et Gabrielle Rocap. « Co-Occurring Synechococcus Ecotypes Occupy Four Major Oceanic Regimes Defined by Temperature, Macronutrients and Iron ». The ISME Journal 10, no 2 (février 2016): 333-45. https://doi.org/10.1038/ismej.2015.115.
2. Farrant, Gregory K., Hugo Doré, Francisco M. Cornejo-Castillo, Frédéric Partensky, Morgane Ratin, Martin Ostrowski, Frances D. Pitt, et al. « Delineating Ecologically Significant Taxonomic Units from Global Patterns of Marine Picocyanobacteria ». Proceedings of the National Academy of Sciences 113, no 24 (14 juin 2016): E3365-74. https://doi.org/10.1073/pnas.1524865113.
